# A deep catalog of protein-coding variation in 985,830 individuals

**DOI:** 10.1101/2023.05.09.539329

**Authors:** Kathie Y. Sun, Xiaodong Bai, Siying Chen, Suying Bao, Manav Kapoor, Chuanyi Zhang, Joshua Backman, Tyler Joseph, Evan Maxwell, George Mitra, Alexander Gorovits, Adam Mansfield, Boris Boutkov, Sujit Gokhale, Lukas Habegger, Anthony Marcketta, Adam Locke, Michael D. Kessler, Deepika Sharma, Jeffrey Staples, Jonas Bovijn, Sahar Gelfman, Alessandro Di Gioia, Veera Rajagopal, Alexander Lopez, Jennifer Rico Varela, Jesus Alegre, Jaime Berumen, Roberto Tapia-Conyer, Pablo Kuri-Morales, Jason Torres, Jonathan Emberson, Rory Collins, Regeneron Genetics Center, RGC-ME Cohort Partners, Michael Cantor, Timothy Thornton, Hyun Min Kang, John Overton, Alan R. Shuldiner, M. Laura Cremona, Mona Nafde, Aris Baras, Goncalo Abecasis, Jonathan Marchini, Jeffrey G. Reid, William Salerno, Suganthi Balasubramanian

## Abstract

Coding variants that have significant impact on function can provide insights into the biology of a gene but are typically rare in the population. Identifying and ascertaining the frequency of such rare variants requires very large sample sizes. Here, we present the largest catalog of human protein-coding variation to date, derived from exome sequencing of 985,830 individuals of diverse ancestry to serve as a rich resource for studying rare coding variants. Individuals of African, Admixed American, East Asian, Middle Eastern, and South Asian ancestry account for 20% of this Exome dataset. Our catalog of variants includes approximately 10.5 million missense (54% novel) and 1.1 million predicted loss-of-function (pLOF) variants (65% novel, 53% observed only once). We identified individuals with rare homozygous pLOF variants in 4,874 genes, and for 1,838 of these this work is the first to document at least one pLOF homozygote. Additional insights from the RGC-ME dataset include 1) improved estimates of selection against heterozygous loss-of-function and identification of 3,459 genes intolerant to loss-of-function, 83 of which were previously assessed as tolerant to loss-of-function and 1,241 that lack disease annotations; 2) identification of regions depleted of missense variation in 457 genes that are tolerant to loss-of-function; 3) functional interpretation for 10,708 variants of unknown or conflicting significance reported in ClinVar as cryptic splice sites using splicing score thresholds based on empirical variant deleteriousness scores derived from RGC-ME; and 4) an observation that approximately 3% of sequenced individuals carry a clinically actionable genetic variant in the ACMG SF 3.1 list of genes. We make this important resource of coding variation available to the public through a variant allele frequency browser. We anticipate that this report and the RGC-ME dataset will serve as a valuable reference for understanding rare coding variation and help advance precision medicine efforts.

---

Exome sequencing has enabled the discovery of rare coding variants which can provide useful insights into gene function^1–4^. Such insights have accelerated the pace of disease gene discovery in both Mendelian studies and common diseases^1,5–7^. In addition to disease allele discovery, exome sequencing has enabled the identification of many protective alleles^8^. Anti-PCSK9 drug therapy is an example of human-genetics-guided drug development based on the observation that loss of PCSK9 function is associated with lower cholesterol levels^9^. Large-scale exome sequencing studies identified rare variants in several genes associated with protective traits, notably, ANGPTL3 for coronary artery disease, GPR75 for obesity, CIDEB for liver disease, MAP3K15 for diabetes, and ANGPTL7 for intraocular pressure^8,10–14^. Thus, exome sequencing efforts have led to the discovery of functionally important rare variants and genes.

Cataloging rare coding variation in genes can aid with the implementation of precision medicine^15,16^. Reference population frequencies are routinely used in understanding the effect of variants on gene function^17^. In genome-wide association studies (GWAS), rare coding variant associations allow unambiguous identification of the causal gene^18^. Furthermore, coding variants can inform the direction of effect and mechanism of action. Here we describe a harmonized collection of ancestrally diverse exonic data obtained from 985,830 individuals, the largest and most diverse data set of its kind^2,19,20^.

## The RGC-ME dataset

### Variation survey

The Regeneron Genetics Center Million Exome dataset contains the genetic variation observed in 985,830 individuals. These data span dozens of collaborations including large biobanks and health systems, disease-specific cohorts, and special population studies. All data were generated by the RGC using a single, harmonized sequencing and informatics protocol. The dataset comprises both outbred and founder populations spanning African (AFR), Admixed American (AMR), European (EUR), East Asian (EAS), Middle Eastern (MEA), and South Asian (SAS) continental ancestries, and includes cohorts with relatively high rates of consanguinity. Approximately 164,000 individuals are of non-EUR ancestry in RGC-ME compared to 34,619 in gnomAD v3.1, 57,290 in gnomAD v2.1, and 90,840 in TOPMed Freeze 8 indicating that RGC-ME has more population diversity than other large scale genetic variation datasets (Fig. 1A)^2,19^.

**Figure 1:**
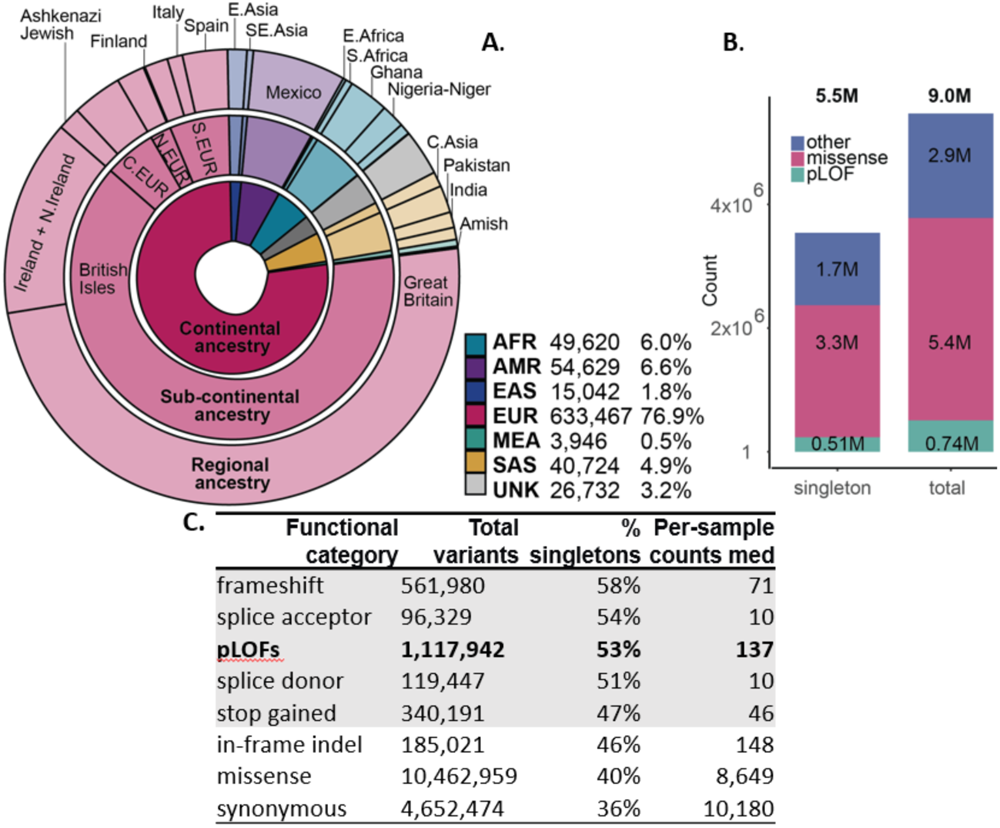
Variant survey and ancestral population counts in RGC-ME. **A.** Summed proportional ancestry (i.e., sum of weighted ancestry probabilities) at continental, sub-continental, and regional levels for 824,159 unrelated samples. All subsequent variant counts and surveys have been performed in this unrelated analysis set **B.** Count of variants unique to RGC-ME that are absent in gnomAD Exomes and TOPMed, broken down by singletons and variant functional category **C.** Variant counts in different functional categories, proportion of singletons, and per-individual median values. All counts are based on variants in the canonical transcript. pLQF includes frameshift, essential splice donor/acceptor (non-UTR) and stop gained categories

We performed a comprehensive survey of genetic variation encompassing single nucleotide variants (SNV) and insertion-deletion (indel) variants. To estimate population allele frequencies, we focused on 824,159 unrelated samples (referred to hereafter as the 824K unrelated set; see Supplementary Table 1 for sample subsets used in various analyses). Sequence analysis identified 32,273,702 SNVs and 2,295,266 indels in autosomal and X chromosomes, of which 36% and 49% are singletons (i.e., only observed in one individual), respectively. Among coding variation in canonical transcripts, 1,117,942 predicted loss of function (pLOF) variants were identified, which include those causing a premature stop, affecting essential splice donor/acceptor sites, or causing frameshifts (Fig. 1C). 53.3% of these pLOF variants are observed as singletons. In addition, 4,652,474 synonymous and 10,462,959 missense variants in canonical transcripts are detected in RGC-ME. 51% of coding variants in canonical transcripts are unique to RGC-ME and absent in other large-scale datasets^2,19^ (Fig. 1B).

Each sample has a median of 137 pLOF, 8,649 missense, and 10,180 synonymous variants (Fig. 1B). AFR individuals have more variants across all functional categories compared to other ancestries (Supplementary Fig. 1), as expected based on the “Out of Africa” model of human population history^21^. Higher variant counts observed in non-EUR may also arise from the use of a single reference sequence as representative of all humans for variant identification.

Mutational saturation of highly methylated CpG sites is almost reached with 1M samples, as previously reported^2^. Of all possible variants in the highly methylated CpG sites, 87.6% synonymous, 84.6% missense, 72% stop gained variants are observed in RGC-ME. Across all mutational contexts, a higher proportion of possible variants are observed in RGC-ME compared to the next largest human exome dataset due to the larger sample size (∼125K and 824K unrelated individuals in gnomAD and RGC-ME, respectively). 21.9% and 8.7% of all possible synonymous and stop-gained variants, respectively, are observed in the RGC-ME compared to 9.3% of synonymous and 3.0% of stop gain variants in a subset of ME with same sample size of 125,000 individuals as gnomAD. Nevertheless, our current sample size is still far from capturing the complete mutational saturation of the human exome (Supplementary Fig. 2).

## Identifying constrained coding regions and genes

### Gene constraint

Population-scale sequencing allows us to quantify pLOF variation in genes, which is key to understanding their role in disease. Several gene constraint metrics (e.g. RVIS, pLI, LOEUF, EvoTol, s_het_) have been developed to estimate tolerance to pLOF variants^22^. Widely used metrics such as pLI^4^ and LOEUF^2^ are derived based on observed and expected pLOF variant counts against background mutation rates but are agnostic to variant allele frequency. A complementary approach to quantifying pLOF depletion considers the cumulative frequency of variants in a gene to estimate a selection coefficient, *s*, on relative fitness loss due to heterozygous pLOF variation^23^ (see Methods). Using this model, we estimated the indispensability of 16,704 protein-coding genes based on the observed number of pLOF variants per gene with cumulative alternate allele frequency (AAF) <0.1% (Supplementary Table 2).

Mean s_het_ in RGC-ME for canonical transcripts is 0.065 [0.037, 0.109] (median s_het_=0.019; Fig. 2A), which is comparable to a mean s_het_ of 0.059 originally computed with the ExAC dataset (n∼60K)^23^. The larger number of samples in our cohort (n∼824k) helps to accurately quantify rare pLOF variants and compute more precise constraint scores than the previously published values based on 60,000 samples. This finding is best illustrated in known haploinsufficient (HI) genes, which are expected to be more constrained and thus have larger s_het_ values relative to all genes (Supplementary Fig. 3). Compared with previously published values from ExAC, s_het_ values for HI genes in RGC-ME are higher on average (Δ*s*_het_= 0.15, p=2.2×10^-21^) and have smaller 95% highest posterior density (HPD) ranges despite those larger means (Δ*s*^_het_ = -0.038, p=1.2×10^-28^). Of the 233 HI genes with s_het_ estimates in both RGC-ME and ExAC, 135 are constrained in both datasets with mean s_het_ >0.07 and lower bound >0.02; 39 genes are classified as constrained only in RGC-ME, while 7 are constrained only in ExAC (Supplementary Table 2). Estimates for all genes were more precise in 824k samples compared to a randomly down-sampled set of 60K samples from RGC-ME (Supplementary Fig. 4A), where mean and median 95% HPD ranges were 6.3 and 4.1-fold larger, respectively. Improving s_het_ estimates is most valuable for shorter genes which were previously difficult to capture at smaller sample sizes. RGC-ME allows more precise estimation of s_het_ for the smallest quantiles of gene coding sequence length (Fig. Supplementary 4B) and derives more informative constraint metrics.

**Figure 2:**
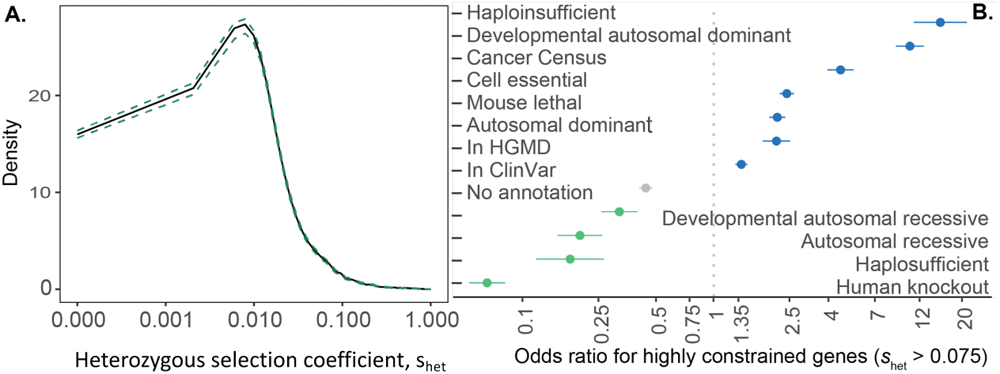
Gene-level constraint estimates representing heterozygous selection coefficients on fitness, s_het_, from RGC-ME. **A.** Mean s_het_ probability density for 16,704 canonical transcripts with 95% confidence intervals calculated with 10,000 bootstrapped samples from means of individual genes. **B.** Odds ratio for genes with s_het_ cutoff > 0.075 to be included in each gene category listed on y-axis.

s_het_ has been shown to be higher in genes associated with Mendelian diseases^23,24^ (Supplementary Fig. 3) and differentiates between groups of genes with different essentiality and disease associations (Fig. 2B). We used s_het_ to identify highly constrained genes in RGC-ME by comparing s_het_ scores in the canonical transcripts of high constraint (haploinsufficient, autosomal dominant in developmental disorders, and lethal in mouse models) and low constraint categories of genes (haplosufficient and genes with rare homozygous pLOF variants; Supplementary Fig. 5). s_het_=0.02 and 0.075 distinguishes genes between these “high” and “low” groups with 81% and 95% precision, respectively. In addition, genes with s_het_ = 0.075 have 81% probability of belonging to the high group relative to the low group. These thresholds serve as cutoffs for mean and lower bound (2.5% HPD) to identify “highly constrained” genes that reflect uncertainty in the mean. Overall, 3,459 highly constrained genes have s_het_ >0.075 and lower bound >0.02. Although 1,241 of these genes lack known human disease associations or lethal mouse knockout phenotypes, they are likely to be of high functional importance. These constrained genes may lack disease associations because loss of even a single copy is incompatible with life or leads to reduced reproductive success without causing clinical disease^25^.

For shorter genes with few expected pLOF mutations^2^, an allele frequency-based approach and larger sample size allow us to estimate constraint metrics not limited by the number of expected pLOF variants. We derived constraint scores for 923 genes underpowered for estimating LOEUF in gnomAD based on ≤5 expected LOFs across 125K exomes. Underpowered genes are significantly shorter with mean coding sequence (CDS) length of 573 bp compared to 1,797 bp for genes with >5 expected LOFs. Most genes with few expected pLOF variants are indeed unconstrained. However, 83 of these are highly constrained with mean s_het_ >0.075 and lower bound >0.02 (Supplementary Fig. 6) and are promising candidates for novel disease gene discovery efforts. 28 have known human disease associations or have been shown to be essential in mouse or cell models. Among these are well-studied genes with known importance in cellular function such as transcription factor TWIST1^26^, DNA and RNA binding protein BANF1^27,28^, and transactivator CITED2.

### Regional constraint

Identifying sub-genic regions intolerant to mutations can reveal functionally important regions otherwise missed when constraint scores are aggregated at the gene level. Models of local coding constraint are powerful tools for identification of protein domains with critical function and for variant prioritization^29–31^. In addition to gene constraint derived from pLOF variation, we also analyzed regional constraint to missense variation using missense tolerance ratio (MTR)^31,32^, defined as the ratio of the observed proportion of missense variants to the expected proportion relative to the number of all possible variants in a defined codon window (see Methods). Using 824K unrelated samples from RGC-ME, we calculated MTR for each amino acid along the CDS within 31 amino acid sliding windows.

Compared with benign missense variants, pathogenic missense variants annotated in ClinVar are highly enriched in the top 1-percentile of MTR constrained regions exome-wide (odds ratio = 18.2; Fig. 3A). Enrichment in pathogenic missense variants persists until the top decile of MTR constrained regions (odds ratio = 1.3). RGC-ME contains a nearly 4-fold larger sample size compared to previous MTR estimates^32^. The increased power from 824k samples improved discrimination between pathogenic and benign variants across MTR percentiles (Fig. 3A) and captured 7% more constrained missense variants (565,127 compared to 604,232) in the top 1 percentile after correcting for false discovery rate (FDR) < 0.1. 0.18% (18,482) of all missense variants (excluding known ClinVar pathogenic variants) observed in RGC-ME have MTR scores in the top 1 percentile and are potentially deleterious. Thus, MTR provides improved interpretation of missense variants and can help prioritize variants in disease gene discovery projects. Top 1 percentile missense constrained regions include important functional regions such as DNA-binding regions, metal-ion binding sites, and active sites (Fig. 3B). Among membrane proteins, transmembrane regions rank higher in MTR constraint regions than cytoplasmic and extracellular domains.

**Figure 3:**
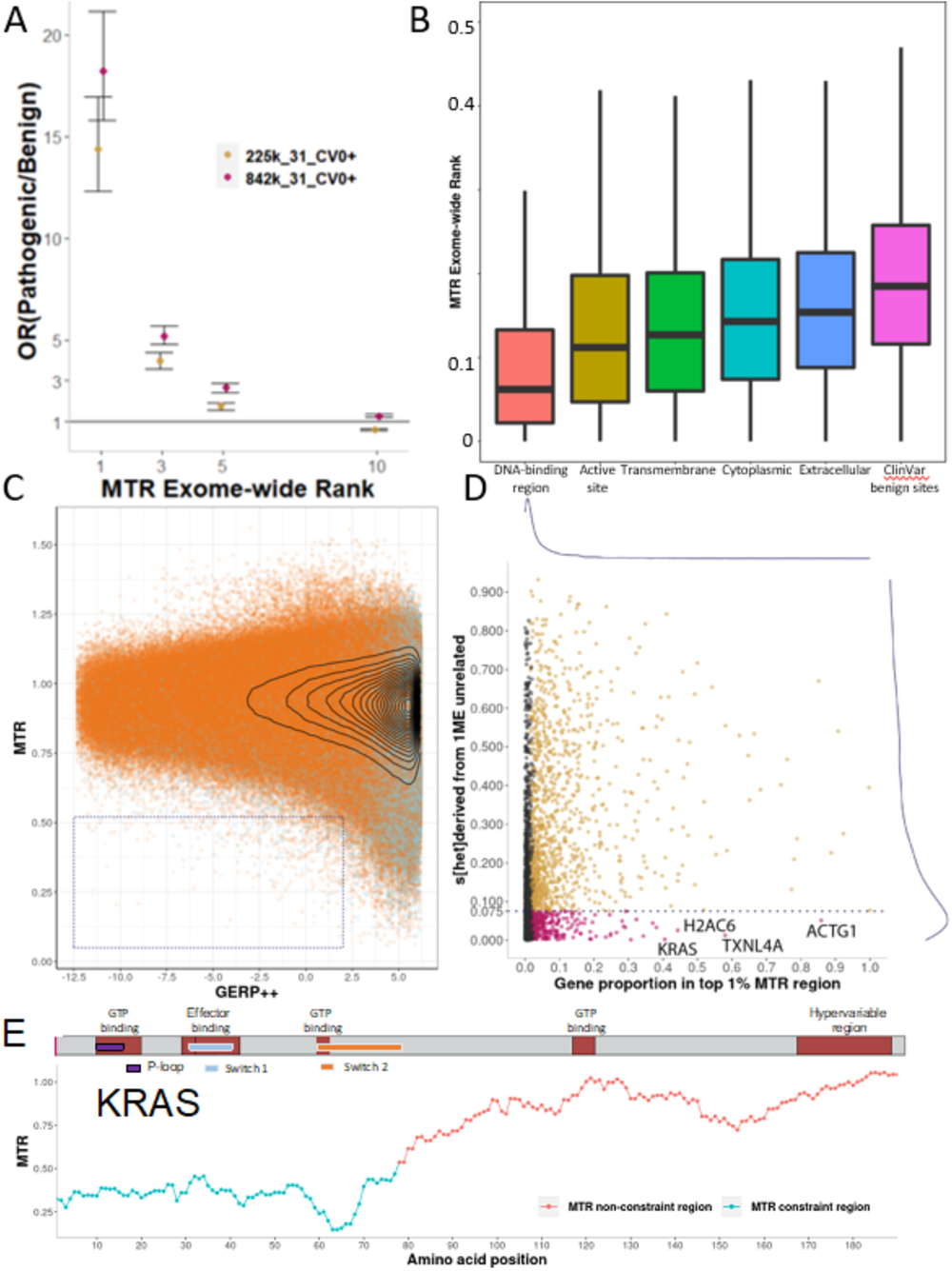
Missense regional constraint captured by MTR. **A.** Odds ratios of ClinVar pathogenic versus benign variants in MTR ranking regions across the whole exome. The pink points represent MTR calculated with a 31 amino-acid sliding window using 824K unrelated samples from RGC-ME and the yellow points represent a random subset of 225K samples. **B.** MTR ranking distribution of different protein functional regions. From left, each category’s distribution of MTR exome-wide ranks was centered at a significantly different location compared to the next category to its right with Wilcoxon rank-sum test (largest p-value =5×10^-10^). **C.** MTR scores of 6.5 million amino acid sites containing missense variants observed from 824K samples against their GERP++ score average on the amino acid site. Cyan dots are overlaid to show sites that are predicted to be deleterious by five missense effect prediction tools. The dotted box highlights sites that are human missense constrained but not cross-species conserved and includes missense variants that have MTR<=0.52 (MTR 1% exome-wide rank) and GERP++ score < 2. **D.** Distribution of the gene proportion located in exome-wide top 1% MTR regions against the heterozygous selection coefficient, s_het_. Genes with significant proportion in most constrained 1% MTR region are colored in orange and red (FDR < 0.1, binomial tests), stratified by LOF constraint (s_het_=0.075). Red dots label genes with missense-specific constrained regions that are LOF-tolerant **E.** MTR track of a cancer oncogene, KRAS, a missense-specific constrained gene, along with the domain structure of the protein. Blue MTR constraint region is defined by top 1% exome-wide MTR rank.

Human intraspecies MTR and inter-species GERP^33^ conservation metrics are uncorrelated for the 6.5 million amino acid changes derived from the observed missense variants in canonical transcripts (Fig. 3C). Although highly conserved sites (GERP++ score ≥2) are enriched in the top MTR constrained sites (MTR value <0.516, 1 percentile MTR exome-wide rank cutoff), linear regression of MTR value and GERP++ score reveals no relationship between the two metrics. 2,643 sites in the top 1 percentile MTR in 818 genes are not cross-species conserved (dotted box, Fig. 3C). The 818 genes with human-specific missense constrained sites are significantly enriched in biological processes of neuronal and immune systems (Supplementary Fig. 7A). Thus, MTR derived from human sequencing data can provide insights into human-specific or recent selection on short evolutionary timescales, complementing cross-species conservation metrics.

To identify highly constrained regions depleted of missense variation, we identified 1,591 genes that have a significant proportion of their coding sequence in the top 1% of MTR (binomial test with π0=0.01, FDR <0.1, Supplementary Table 4). We further compared the genes that are highly missense-constrained to the LOF-constraint metric, s_het_ (Fig. 3D). 292 of the 1,591 genes are not LOF-constrained, i.e., s_het_ <0.075. Additionally, 165 missense-specific constrained genes do not have s_het_ estimates due to lack of pLOF variants or data quality. These 457 genes with missense-specific constrained regions have significantly shorter CDS length than 641 LOF-specific constrained genes (p= 8.38×10^-30^, Supplementary Fig. 7B). pLOF variants are under the greatest selection and it is not possible to obtain region-level LOF constraint due to the paucity of pLOF variation. MTR serves as a complementary lens for identifying functionally important regions at a higher resolution than gene level LOF constraint. Moreover, MTR also identifies functional regions in genes that are tolerant to LOF variation. For example, KRAS, a well-known oncogene, is LOF-tolerant to (s_het_ =0.002, LOEUF=1.24); however, the first 77 amino acids (40%) of the protein sequence are ranked in top 1% MTR constrained regions (Fig. 3E). This region includes the P-loop, Switch 1 and Switch 2 functional domains which form critical binding interfaces for effector proteins^34^ and highlights the importance of regional constraint metrics.

### Understanding human knockouts

RGC-ME individuals span six continental ancestries and include founder populations and cohorts with high rates of consanguinity, contributing to the most comprehensive collection of homozygous loss-of-function variation compiled to date^35–37^. Overall, we identified 4,874 genes with rare, homozygous pLOF variants, providing an opportunity to understand the effect of gene function directly from phenotypic characterization of individuals harboring such variants, effectively, naturally occurring “human knockouts.” We assume pLOF variants that affect both copies of an allele inactivate the gene. Homozygous pLOFs in RGC-ME are generally rare; 65% are found in only one participant and 41% of gene knockouts have one variant (Fig. 4A). The 4,874 putative gene knockouts (pKO) comprise 9,135 rare (AAF<1%), homozygous pLOF variants identified in 49,971 individuals from RGC-ME data (5% of 985,830 samples). 1,838 of the pKOs have not been previously reported (Supplementary Table 5). As expected, cohorts with higher rates of consanguinity^35,36,38^ are enriched in pKOs compared to outbred populations despite smaller sample sizes (Fig. 4B-C, Supplementary Table 6).

**Figure 4:**
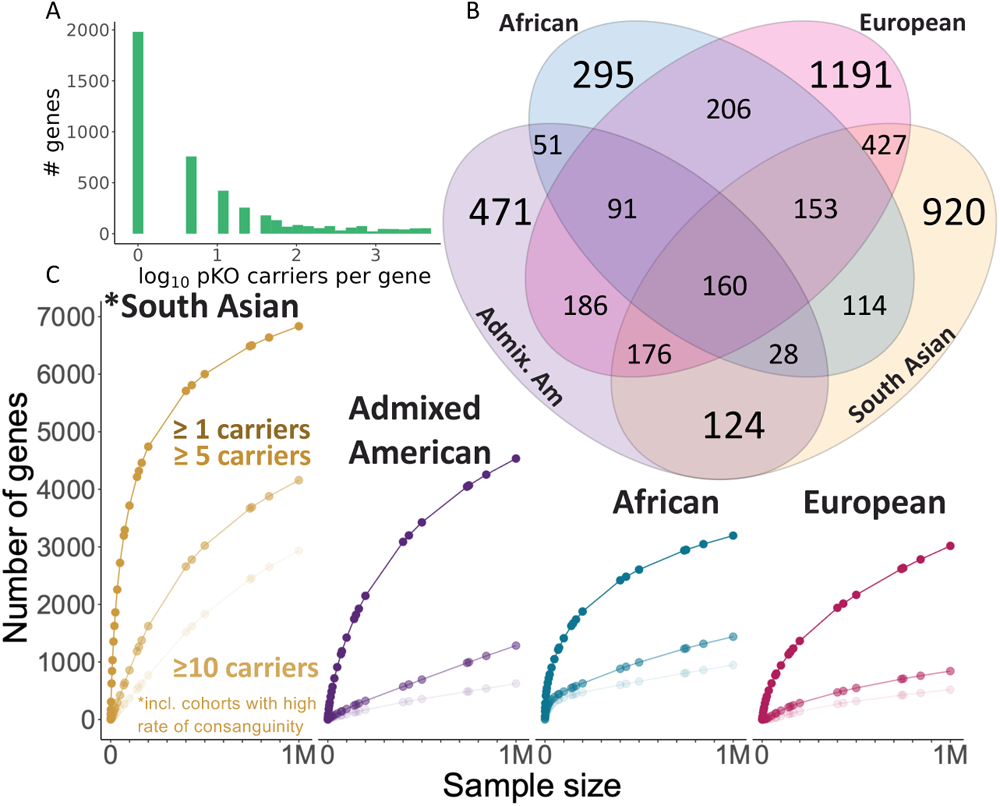
Rare homozygous pLOF variants and “human knockouts” in RGC-ME. **A.** Distribution of the number of individuals per gene knockout on the log10 scale **B.** Breakdown of the number of putative gene KOs observed in RGC-ME by ancestry. **C.** Projected accrual of putative gene KOs at hypothetical cohort sizes for each ancestral group in 1.01M related individuals. Curves reflect accrual of the expected number of genes with at least 1, 5, and 10 carriers, respectively, of a homozygous variant.

pKOs are significantly less constrained than expected. Only 2.3% of pKOs have s_het_ >0.075 compared to 20.9% of all human genes (p=1×10^-307^) and 57% of pKOs are in the lowest decile of s_het_ scores exome-wide (s_het_ < 0.008). A caveat is that s_het_, like most gene-specific measures of constraint, is designed to capture selective pressures on heterozygous pLOF variant carriers^39^. Although genes harboring homozygous pLOF variants are under less selective pressure, current sample sizes are inadequate^40^ to directly compute selection on homozygous variation.

pKOs are overrepresented in drug and xenobiotic metabolism pathways. 34 of the 57 human cytochrome P450 (CYP) genes have rare or common homozygous pLOF variants. We observe notable allele frequency differences across population among variants in CYP genes. For example, AMR and SAS individuals have higher pLOF frequencies in CYP2C19, whereas AFR individuals have higher pLOF frequencies in CYP2C9 and CYP2A6 (Supplementary Fig. 8). Polymorphisms in EUR and AFR individuals in CYP2D6 and in EAS individuals in CYP2C19 are associated with differential metabolism of widely used drugs, such as beta blockers, antidepressants, and opioids^41–44^. Genetic variation within these CYP genes may be determinants of poor or rapid drug metabolism and using pLOF variants as genetic proxies for therapeutic gene-knockout can help to further precision genetic medicine efforts. Other well represented gene families include solute carriers (SLC: 128 of the 362 have knockouts), olfactory receptors (OR: 58 out of 390), and ATP-binding cassette transporter genes (ABC: 23 out of 49). The presence of human knockouts in these gene families suggests that there may be functional redundancy between homologous genes.

Characterizing genes that tolerate loss of both alleles may identify drug targets that can be knocked out with minimal side effects^35^. Drug targets with homozygous pLOF variants in humans are more likely to progress from phase I trials to approval^36^. Of 1,175 inhibitory preclinical targets listed in the Drug Repurposing Hub, 187 (16%) have at least one individual with a rare homozygous pLOF variant in RGC-ME^45^. In-depth phenotyping of human knockouts can help researchers better understand the efficacy and side-effect profiles of these potential drug targets. Furthermore, human knockouts provide a way to understand the consequences of life-long deficiency of a gene^46^. Thus, meticulously cataloguing a “human knockout roadmap” is useful for drug development efforts.

### Identifying variants with differential frequency across populations

Adaptation to constant selective pressures such as diet, climate, and pathogen load have largely contributed to global human diversity. The breadth of continental ancestries represented in RGC-ME provides an opportunity to identify coding variants under the influence of directional selection and/or drift, resulting in higher differentiation of beneficial alleles in one population compared to others. Several loci important for human adaptations have been discovered, such as associations with adaptation to higher altitude^47^, protection from malaria^48,49^ and black death^50^, and other immune and metabolic traits^51^. In many instances, the higher frequency of differentiated alleles in selected populations compared to well-studied European populations results in improved power in association analysis and can aid in identification of novel associations of genetic variation to medically relevant phenotypes^52^. For example, functional variants within alcohol metabolism genes and their role in protection to alcohol consumption was discovered due to higher differentiation of these variants in the East Asians^53–55^.

To identify such highly differentiated variants, we calculated the degree of population differentiation for ∼35 million variants using the Fixation Index (FST) within each major population (AFR, AMR, EAS, EUR, SAS; Supplementary Table 7.1 and 7.2). We observed high FST values (FST > 0.15) for multiple variants in both known and novel loci, which represent putatively positively selected loci in each population (Supplementary Fig. 9; Supplementary Table 7.1). A large proportion of variants in AFR and EAS show a higher degree of differentiation than AMR and SAS when compared against EUR allele frequency. Similar results are seen when each ancestry is compared against the pooled allele frequency data across all ethnicities.

Many high FST variants in AFR and EAS have low allele frequency in EUR (Fig. 5). Several were associated with traits ranging from morphological (skin/hair pigmentation), blood related (hemoglobin levels, reticulocyte counts), and complex disorders like type 2 diabetes mellitus (T2D) in summary statistics from Biobank of Japan and UK-Biobank^5^ (Supplementary Table 7.3 and 7.4). Interestingly, T2D, metabolic disorder risks and immune system disorders has been previously hypothesized to be a target of natural selection in humans because of adaptations to metabolism, energy production, and pathogens^51,56–58^. Some of the notable association signals of genes with variants having high Fst identified in Biobank of Japan data that fit this hypothesis include *PAX4* (p=6.3 x 10^-90^; T2D), *GP2* (p=1.33×10^-14^; T2D), *GLP1R* (p= 6.10×10^-14^; T2D), *ATXN2* (p=1.09×10^-11^; chronic obstructive pulmonary disease), 6p21.32 (*BTNL2*, *NOTCH4*; p<2.89×10^-54^; rheumatoid arthritis) and *NLRP10* (p=2.39×10^-35^; atopic dermatitis) (Supplementary Table 7.4). We also observed significant association of high-Fst variants in *IFRD2* (p=2.72 x 10^-21^; reticulocyte count), *MFSD2B* (p=1.75 x 10^-12^; HbA1C), *USP40* (p=1.05 x 10^-10^; total bilirubin) in UKB-Africans summary statistics. Notably, these variants have low frequency in EUR and thus have low power to be identified in EUR cohorts. Variants that showed high FST (FST > 0.05) were significantly enriched for phenotypic categories related to coronary artery disease (p= 8.15 x 10^-5^) and metabolic disorders (p= 1.01 x 10^-4^) in the GWAS catalog.

**Figure 5:**
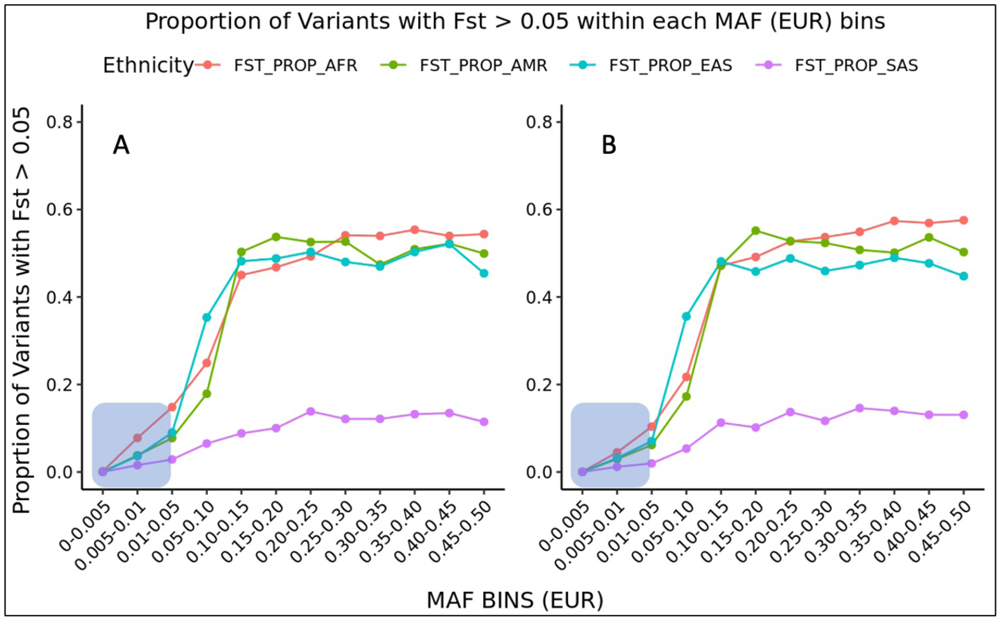
Fst distributions across allele frequency and functional classes. Proportion of high FST (>0.05) variants by allele frequency in Europeans for (A) synonymous variants and (B) missense variants. Several European rare/low-frequency exonic variants (shaded blue area) are more differentiated in Africans, Admixed Americans, and East Asians compared to South Asians.

As previously reported, we also found limited overlap between highly differentiated variants (variants with high Fst) and phenotype-associated variants in SAS, AFR, and AMR^56,59^. One reason might be that high impact and large effect size variants are quickly eliminated or maintained at lower frequency in the population. We indeed observe enrichment of coding variants at lower FST bins in RGC-ME (Supplementary Fig. 9.2). The limited phenotypes characterized in GWAS studies as well as the small case numbers for several traits in AFR, AMR, and SAS are known impediments to understanding signatures of positive selection^59^. Nonetheless, as GWAS for additional traits become available, our extensive catalog of novel differentiated variants will continue to aid in the understanding of evolutionarily interesting loci.

### Interpretation and annotation of variants that affect splicing

Large scale genetic cohorts capture rare variation which can be used to understand selection and functional importance of variants. We used RGC-ME to characterize deleterious variants, which are expected to have lower allele frequencies than neutral variants due to negative selection. We can infer the degree of selection between different functional classes of variation by comparing the proportion of singletons (variants observed only once) in each class. Accounting for both background mutation rate and methylation level^60^, we computed the deleteriousness of variants using an updated mutability adjusted proportion of singletons (MAPS) metric^4^. As expected^2,4^, pLOF variants had the highest MAPS scores, followed by missense, synonymous, and non-coding variants, respectively (Fig. 6A). To systematically assess the deleteriousness of variants that may affect splicing, we used splice prediction scores from SpliceAI and MMsplice^61,62^ to group variants observed in RGC-ME into predicted score bins and compared the MAPS scores of variants in each bin (Supplementary Fig. 10). We identified deleterious variants that affect splicing as those with a minimum MAPS score equivalent to missense variants predicted to be deleterious by five different prediction models, i.e., 5/5 missense variants. This corresponded to a SpliceAI score threshold of 0.37 and MMSplice score threshold of 0.97.

**Figure 6:**
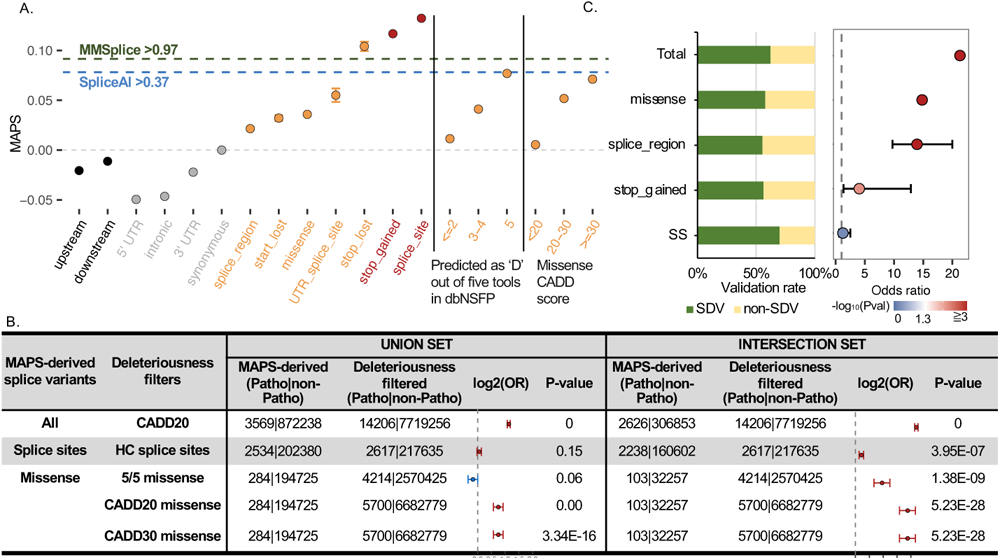
Identification of variants that are predicted to affect splicing. A. The mutability-adjusted proportion of singletons (MAPS) across different functional categories. Error bars represent standard error of the mean of the proportion of singletons. The blue and green dashed lines represent the SpliceAI and MMSplice score thresholds respectively for variants that have a MAPS score equal to that of missense 5/5 (predicted deleterious by 5 algorithms) variants. Variants with spliceAI score >= 0.37 or MMSplice score >= 0.97 are predicted to be deleterious splicing-affecting variants B. Enrichment of ClinVar pathogenic variants in predicted splice-affecting variants compared with corresponding variant sets filtered by either LOFTEE, 5/5 missense deleteriousness models, or CADD20 C. Empirical validation of MAPS predicted splice-affecting variants **Left panel:** fraction of predicted splice-affecting variants (intersection set) validated as splice disrupting variants (SDVs) in any of the three splice reporter assays **Right panel:** enrichment of predicted splice-affecting variants in SDVs compared to non-SDVs

We identified 293,332 coding variants that are predicted to affect splicing in RGC-ME based on the intersection set (i.e., both by SpliceAI and MMSplice based on MAPS-derived splicing thresholds; Supplementary Fig 11A). 42.8% (125,556) of all predicted splice-affecting variants are non-splice sites (non-SS). SS and variants within the splice region comprised the largest category of variants predicted to affect splicing. SpliceAI and MMSplice identified 75% of LOFTEE high confidence (HC) SS and ∼10% of variants within splice regions as splice affecting (Supplementary Fig. 11A-B). Both SpliceAI and MMSplice also identified ∼50% of LOFTEE low confidence (LC) SS as deleterious splice-affecting variants. The impact of non-SS variants on alternative splicing is often underestimated. For example, missense and synonymous variants accounted for 36% and 18% of all predicted splice-affecting variants identified by MAPS-derived spliceAI and MMSplice thresholds, respectively (Supplementary Fig. 11A-B), indicating that non-SS variants have a non-negligible impact on splicing.

MAPS-derived variants predicted to affect splicing are enriched in well-supported ClinVar pathogenic variants (STAR ≥2) compared to other widely used variant deleteriousness metrics. Variants in both the union (i.e., identified by either SpliceAI or MMSplice) and intersection sets of predicted splice-affecting variants are significantly enriched in ClinVar pathogenic variants compared to variants with CADD>20. SS in the intersection set are significantly enriched for pathogenic variants than LOFTEE HC SS, indicating that LOFTEE may remove some true deleterious splice sites. Moreover, missense variants in the intersection set were significantly enriched in pathogenic variants compared to 5/5 missense variants (variants predicted to be deleterious by five out of five prediction algorithms in dbNSFP) and variants with high CADD scores (*i.e.,* CADD>20 and CADD>30; Fig. 6B). Non-SS cryptic splice variants may thus be important in human disorders, and these results suggest that MAPS-derived thresholds for predicted splice scores can identify deleterious splicing variants.

MAPS-derived splice score thresholds also detected experimentally verified variants that affect splicing. We compiled a dataset of 36,067 SNVs from three high throughput splicing reporter assays: MapSy, Vex-Seq, and MFASS^63–65^, 2,806 of which were reported as large-effect splice disrupting variants (SDVs; Supplementary Table S8). 585 SDVs (out of 2,558 SDVs with either spliceAI or MMSplice prediction scores) were identified using the MAPS-derived splice prediction score threshold. MAPS-derived thresholds for SpliceAI and MMSplice (intersection set) identified around 70% of SS, 56% of stop gain variants, and more than half of missense, and synonymous SDVs (Fig. 6C left panel). Importantly, predicted splice-affecting variants identified by MAPS-derived thresholds were significantly enriched in SDVs compared to non-SDVs in all but SS category, whose odds ratio is also higher than 1 (Fig. 6C right panel). Similar results were obtained using the union set (Supplementary Fig. 11C).

### Interpretation of rare variants for clinical diagnostics

To understand the prevalence of disease alleles in the general population, we identified well-supported ClinVar^66^ pathogenic variants (STAR ≥2) in the 824K unrelated RGC-ME samples. 20,412 (40.8%) ClinVar2+ pathogenic variants were observed in RGC-ME, of which 20,331 (99.6%) have AAF < 0.1% and 3,638 (17.8%) were observed only once. In comparison, only 20% (9,915) and 30% (14,862) of pathogenic variants were observed in ExAC (n≈60K) and gnomAD (n≈125K) exomes respectively, indicating that the larger sample size of RGC-ME allows us to identify many previously unobserved rare pathogenic variants (Supplementary Fig. 12). Each RGC-ME participant harbors on average 1.6 pathogenic variants, with 0.33 variants in 570 autosomal dominant genes and 1.3 variants in 1,299 autosomal recessive genes based on inheritance mode annotations from OMIM.

The American College of Medical Genetics identified a set of genes (ACMG SF V3.1) with clinically actionable variants known to predispose individuals to diseases and with medical interventions available to improve mortality and morbidity^67^. Among the 824K unrelated individuals, 22,846 (2.77%) have at least one high confidence pathogenic (P) missense or pLOF variant reported in ClinVar for 72 of the 76 autosomal genes on the list (Supplementary Table 9). As expected, two of the most prevalent pathogenic variations were the HFE p.Cys282Tyr allele, with 3,344 homozygous individuals (enriched in EUR, nEUR-hom=3,223 and AAFEUR=13.8%), and the TTR p.Val142Ile allele with total allele count (AC) of 2,085 (majority AFR carriers, nAFR=1,676 and AAF=3.4%). We also tallied carriers of likely-pathogenic (LP) pLOF variants in 44 genes where truncation is known to lead to disease. 2,357 (0.3%) individuals in RGC-ME carry 1,407 LP variants across 40 of 44 of these genes. In total 3.06% of RGC-ME are carriers of P or LP variants. Excluding individuals with high frequency pathogenic variants in the HFE (Cys282Tyr) and TTR (Val142Ile) genes, 2.38% of RGC-ME carry an actionable variant (Supplementary Table 9). Pathogenic variant findings are expected to be rare in large-scale general population efforts and 39% and 80% P and LP variants, respectively, are singletons.

Since RGC-ME has the largest-to-date uniformly processed exome data from continental ancestries other than EUR, we interrogated the prevalence of all pathogenic variants in ClinVar across three ancestral populations – AFR, AMR and SAS – in addition to EUR. Ancestral groups were sub-sampled 5 times for equal sample sizes of 29,521, matched to the smallest ancestry group, SAS. We tallied variants across both known and unknown pathogenic classes (“unknown significance” and “conflicting interpretations of pathogenicity” comprise the latter) annotated by ClinVar, broadening our analysis beyond high confidence pathogenic variants. Understanding variants of unknown significance (VUS) is currently a bottleneck in interpretation of variation in clinically relevant genes and a challenge in clinical management^15^.

On average, we observed 11,500 high confidence pathogenic coding (missense and pLOF) variants in each random sub-sample of ∼30k individuals per ancestry (∼120k individuals total across 4 ancestries) and we observed a higher number among EUR. 57% of such variants observed per sub-sample on average were found in EUR samples compared to 44%, 31% and 34% in AFR, AMR, and SAS, respectively, indicating the range of known pathogenic allelic representation in ClinVar (s.d. ± 0.07-0.35% for EUR, AFR, and AMR). On average, 2,727 pathogenic variants were unique to EUR per random down-sample, compared to 1,575, 1,342, and 1,107 unique to AFR, SAS, and AMR, respectively, corresponding to a 73-146% increase in variants private to EUR. Nevertheless, sampling diverse populations increases the likelihood of capturing alleles that are rarer or absent in Europeans: about 35% of unique pathogenic coding variants per sub-sample were only observed in non-EUR samples.

Next, we looked at the allele frequency distributions of VUS across various ancestries. Overall, VUS are at higher allele frequencies (average AAF=3.75×10^-4^) than pathogenic variants (average AAF=9.75×10^-5^). Only 11 pathogenic coding variants on average in AFR, 5 in AMR, and 4 in SAS are >100x more common in these respective ancestries than EUR. In contrast, 1.2% of VUS found in non-EUR populations (4,009 variants, sd=30.2) have ≥100-fold AAF compared to EUR on average (Supplementary Fig. 13A). In AFR, over 500 and 700 of these variants have > 5% and 1% AAF, respectively (Supplementary Fig. 13B). Rare variants that are common in at least one population should be thoroughly evaluated for pathogenicity. While there are documented examples of disease alleles with elevated allele frequencies in population-specific founder variants,^68–71^ detailed phenotypic and functional characterization of variants is critical for variant curation efforts.

Although VUS have less empirical evidence for pathogenicity, they comprise the bulk of Clinvar with >1 million variants and truly deleterious variants are among their ranks. Previously, we identified missense constrained regions of the exome using MTR in RGC-ME. 6,117 VUS (0.58% of missense VUS) in ClinVar map to top 1% MTR constrained regions and 24,511 (12%) VUS map to top 5 regions (Supplementary Table 10). These variants are in regions of greater human constraint and may be best classified as likely pathogenic. In addition, we extended our previous use of the MAPS-derived splicing score thresholds to identify potentially deleterious variants that may affect alternative splicing in ClinVar. In the unrelated RGC-ME population, there are 603,727 missense and 30,999 synonymous variants annotated as CI or VUS. Among them, 4.1% (1,269) of synonymous and 1.6% (9,439 variants) of missense CI/VUS can be annotated as cryptic splice variants based on the union set. While these numbers represent a small proportion of the ClinVar variants lacking unambiguous pathogenic annotations, they include over 10,000 candidate variants that can be functionally assayed for splicing changes and downstream interpretation of their effects. Characterizing VUS in constrained regions or characterizing them as splice affecting will enable improved annotation and interpretation for clinical diagnostics.

## Discussion

The RGC-ME dataset builds upon previous efforts such as gnomAD to characterize human genetic variation with a harmonized catalog of ∼35 million high quality variants, representing the largest collection of rare exonic variants and sequenced exomes to date. Our dataset comprising 985,830 exomes is publicly available (https://rgc-research.regeneron.com/me/home) featuring variant frequency data for globally representative ancestral groups. Large-scale public variant repositories have been instrumental in facilitating genomic discoveries. Here, we describe characteristics of the variant-and gene-level data in RGC-ME to preview the value of this public resource.

The sample size of RGC-ME, ancestral diversity, and inclusion of cohorts with high rates of consanguinity in the dataset adds over 1,800 new genes with rare homozygous pLOFs to the human knockout catalog. Studying KOs affords the opportunity to identify drug targets with improved safety profiles by deeper phenotyping of such individuals. Homozygous pLOF variants arise in genes that can tolerate mutations which may indicate lower biological essentiality, greater diversity across populations, or redundancy among genes in large families. For example, the cytochrome P450 superfamily, well-represented among human knockouts in RGC-ME, has differential allele frequencies across populations and pharmaceutical relevance. Understanding individual differences in drug response will be critical as medicine becomes increasingly personalized.

The breadth of RGC-ME dataset helps identify genes on both extremes of the spectrum, from putative gene KOs tolerant of homozygous truncating mutations to constrained genes with few pLOF variants. Generating constraint metrics based on rare pLOF variant frequencies and incorporating data from a larger sample size improves our power to detect constraint in genes with few expected pLOF variants. We demonstrated improved predictions in the most underpowered set of protein-coding genes (≤5 expected LOFs) and detect constraint in short genes of known cellular importance. Although uncertainty persists in these estimates, we can confidently call over 80 of these genes as intolerant to pLOF variants, some of which are critical to cellular function such as the transcription factor TWIST1. RGC-ME also provides unprecedented power to measure missense constraint across sub-regions in the whole exome, differentiating between pathogenic and benign variants better than previous estimates. Larger sample sizes capture missense constrained regions more effectively by enriching pathogenic risk in MTR constrained sites. MTR captures regional constraint across >450 genes where LOF constraint is underpowered or undetectable, or where gene-level metrics are insufficient to identify sub-regional constraint.

Cataloging variation at scale provides an opportunity to observe rare pathogenic variation. We observed fewer ClinVar variants among samples of AFR, AMR and SAS ancestries compared with EUR. In sub-samples of equal sizes across ancestries, 23% of high confidence pathogenic ClinVar variants observed in RGC-ME were only present in EUR. This suggests that existing medical variation data may be under-representative of diverse populations, particularly since AFR harbor more rare variants on average per individual. Recruitment of diverse cohorts to identify and characterize novel pathogenic variants will help address this ascertainment bias, along with increased access to genetic medicine. Most disease-associated genes and variants remain uncharacterized despite rich annotation resources and clinical cohorts. VUS are difficult to classify for several reasons, including the lack of population-based statistical evidence, functional evidence, and different professional evaluation criteria. Characterizing variants from diverse ancestries and their relative frequencies will improve pathogenic variant annotations by including more representative allele frequency distributions^72^.

One avenue to further annotate disease relevant variants is to understand the impact of non-splice site variants that affect splicing. Identifying these cryptic splice variants, however, is challenging. Prediction tools can identify splice-affecting variants, but the deleteriousness of variants based on splicing prediction score thresholds has not been systematically assessed. Using a data-driven approach to quantify the degree of natural selection against different functional variant categories, we determined appropriate splicing prediction thresholds for state-of-art models (i.e., SpliceAI and MMSplice) in capturing deleterious splice variants. We identified over 7,000 putative splice disrupters among ClinVar VUS annotated as missense or synonymous variants. Refining variant annotation and providing additional context for causative mutations can illuminate disease mechanisms and next steps for functional prioritization and clinical utility.

Reference population databases are widely used to interpret gene and variant biological functions^17,73^. Here, we expand upon the proven utility of large-scale variant datasets with an even deeper catalog capturing greater genetic diversity. We provide a public browser (https://rgc-research.regeneron.com/me/home) containing variant and allele frequency data on a fine ancestry level to advance genomic research. The catalog of coding variation in RGC-ME will be an invaluable resource for rare variant interpretation and is a step forward towards the realization of precision medicine.

## Methods

We aggregated high quality whole-exome sequencing data from 985,918 individuals after removing samples in a rigorous quality control process based on a variety of sequencing metrics. To ensure this sample set characterizes genetic variation representative of the general population, we excluded available samples from Mendelian cohorts, samples with neurodevelopmental disorders, and samples with use-restrictions inconsistent with the analysis presented here.

### Sample preparation and sequencing

Genomic DNA libraries were created by enzymatically shearing high molecular weight genomic DNA to a mean fragment size of 200 base pairs. Multiplexity of exome capture and sequencing was achieved by adding unique asymmetric 10-bp barcodes to the DNA fragments of single samples during library amplifications. Equal molar amounts of DNA samples were pooled for exome capture using a slightly modified version probe library of xGen exome research panel from Integrated DNA Technology (IDT). After PCR amplification and quantification of the captured DNA, samples were multiplexed and loaded to Illumina sequencing machines for sequencing to generate 75 base pair paired end reads. The samples in this study were sequenced using the Illumina sequencing machines including HiSeq 2500, and NovaSeq 6000 with S2 or S4 flow cells.

### Read mapping and variant calling

Sequencing reads in FASTQ format were generated from Illumina image data using bcl2fastq program (Illumina). Following the OQFE (original quality functional equivalent) protocol^74^, sequence reads were mapped to GRCh38 references using BWA MEM^75^ in an alt-aware manner, read duplicates were marked, and additional per-read tags were added. Single nucleotide variations (SNV) and short insertion and deletions (indels) were identified using a Parabricks accelerated version of DeepVariant v0.10 with a custom WES model and reported in per-sample genome VCF (gVCF)^76^. These gVCFs were aggregated with GLnexus v1.4.3^77^ into joint-genotyped multi-sample project-level VCF (pVCF), which was converted to bed/bim/fam format using PLINK 1.9^78^ for downstream analyses.

### Quality control (QC) of dataset

We implemented a support vector machine-based QC protocol to filter likely artifactual variants as previously described^5,38^. Positive controls were defined as: (i) genotype calls with ≥ 99% concordance between array and exome sequencing data; (ii) transmitted singletons; and (iii) an external set of likely “high quality” variants defined from 1000 genomes phase 1 high-confidence SNPs and Mills and 1000 genomes gold-standard INDELs, further restricted to the intersection between variants that pass QC in TOPMED Freeze 8 and gnomAD v3.1.2 genomes. Negative controls were defined as: (i) mendelian inconsistent variants (where # of mendel errors (ME) ≥ 3 and ME – allele count (AC) ratio ≥ 0.01); (ii) discordant genotype calls in genomic duplicates (where # of discordant calls (DC) ≥ 3 and DC – AC ratio ≥ 0.05); and (iii) intersection of gnomAD v3.1.2 fail variants with TOPMED Freeze 8 mendelian or duplicate discordant variants. Prior to model training, the control set of variants were subset to exome target capture regions only, binned by allele frequency (AF), and then randomly sampled such that an equal number of variants were retained in the positive and negative labels. The model was then trained on up to 36 available site quality metrics, including, for example, the median value for allele balance in heterozygote calls and whether a variant was split from a multi-allelic site. Even and odd chromosomes were then split into train and test sets, respectively. We performed a grid search with 5-fold cross-validation on the training set to identify the hyperparameters that return the highest accuracy during cross-validation, which are then applied to the test set to confirm accuracy (Precision=0.92, Recall=0.98, F1=0.95, AUC=0.98). Further, we find that for unobserved labels (variants that were withheld from model training and testing due to sample matching within AF bins), 97% of true positives are predicted to pass and 92% of true negatives are predicted to fail. This approach identified as low-quality a total of 2,838,213 (11%) variants in exome target capture regions.

### Variant annotation

Variants were annotated using Variant Effect Predictor, VEP^79^, v100.4 based on Ensembl^80^ release 100 human protein-coding transcript models. The annotation is primarily based on protein-coding transcripts that have a defined start and stop codon. Incomplete transcripts are not included apart from a small number that have a MANE^81^ annotation.

We used a single consequence per variant based on its annotation on the canonical transcript for all the analyses described in the main text. One canonical transcript per gene is defined using a combination of MANE^81^, APPRIS^82^ and “Ensembl canonical” tags. “Ensembl canonical” transcript was defined using the following hierarchy: 1. Longest CCDS translation with no stop codons. 2. If no (1), choose the longest Ensembl/Havana merged translation with no stop codons. 3. If no (2), choose the longest translation with no stop codons.

The definition for “Ensembl canonical transcript” was obtained from http://dec2017.archive.ensembl.org/info/website/glossary.html

MANE annotation is given the highest priority, followed by APPRIS. When no MANE or APPRIS annotation tags are available for a gene, the Ensembl canonical transcript definition is used. Transcripts with a “MANE Select v0.91” tag were used to identify the canonical transcript of a gene. APPRIS and Ensembl tags were obtained from protein-coding transcripts derived from the human Ensembl Release 100 build.

LOFTEE, an Ensembl-VEP plug-in for pLOF variants was separately run on VCF v4.2 files containing all observed pLOF variants in RGC-ME (stop gained, splice acceptor/donor, frameshift). Analyses with pLOF variants were restricted to High Confidence (HC) variants unless otherwise specified.

### TOPMed imputation

Separately for each individual input cohort, array data, generated from Illumina Omniexpress or Global Screening Array (GSA) v1 and GSA v2 beadchips according to manufacturer’s recommendations, was filtered (MAF > 1%, HWE p-value > 1e-15, site-level missingness < 1%) and split into chromosome and sample batches using PLINK2. Each batch was submitted as a job on the TOPMed Imputation Server (https://imputation.biodatacatalyst.nhlbi.nih.gov). On the server, the dataset was phased using eagle v2.4 and imputed using MINIMAC4 against the 97,256 deeply sequenced genomes in the TOPMed reference panel. When complete, the imputed data in VCF files were retrieved from the server and merged sample-wise using bcftools in 5MB genomic regions. Variants were then converted from VCF to PGEN format by reading the HDS field.

### Fine-scale ancestry (FSA) assignment

To estimate genetic ancestry in RGC-ME we created a reference set of 11,354 individuals from 95 populations by combining several fully public, approved access and internal array genotyping and whole genome sequence datasets^83–90^. We then calculated the proportion of haplotype sharing of each RGC-ME sample to the 95 populations in the reference set utilizing a haplotype sharing model^91^ and applied to the RGC-ME TOPMed imputed array dataset. Individuals were assigned to a population if they had greater than 50% ancestry from that population. Populations were also grouped into 6 sub-continental groups. Individuals with greater than 50% ancestry from a group were then hard assigned to that group: African (AFR) N = 64,600, Admixed American (AMR) N = 85,726, East Asian (EAS) N = 7,565, European (EUR) N = 872,756, Middle Eastern (MEA) N = 3,130 and South Asian (SAS) N = 48,076. A total of 53,260 individuals were unassigned using this approach.

### Amish allele frequency estimation

The Amish could not be assigned using the above FSA method because the reference set did not include any Amish individuals. For estimation of allele frequency of variants in the Amish population, we performed PCA on 1,963 samples from cohorts with predominantly Amish individuals and the same number of samples randomly selected from UKB450K cohort. We further used K-means clustering on PC1 values, which captures the most variation across samples (Supplementary Fig. 14), to distinguish between Amish and non-Amish EUR samples. All non-Amish EUR samples were grouped in one cluster (blue cluster), and 1,225 (out of 1,963) initial Amish samples were grouped in another (red cluster, Supplementary Fig. 14). Allele frequencies for the Amish were estimated from the sequence data of the 1,225 samples

### Genetic relatedness analysis

To estimate the portion of identical-by-descent (IBD) genomic regions shared between pairs of individuals in our study, we first obtained a set of high-quality common SNPs from the exome variant set by excluding SNPs with MAF < 10% and genotype missingness > 5% and all indels. Further, variants with abnormal het rates based on the expected (exp) vs observed (obs) het calculations based on empirically determined cutoffs (obs - exp > 0.01 or exp - obs > 0.1) were excluded from further analysis. The asymmetry in the cutoff values was selected to account for the Wahlund effect. IBD estimates were calculated among individuals within the same ancestral superclass that was determined by projecting each sample onto reference principal components calculated from the HapMap3 reference panel using PLINK with a minimum PI_HAT cutoff of 0.1875 to capture out to second-degree relationships, which generates an ancestry-version IBD estimates. A separate IBD estimation was calculated among all individuals using a minimum PI_HAT cutoff of 0.3 to identify the first-degree relationships among all samples to generate “first-degree family networks”, which are connected components of individuals (nodes) and first-degree relationships (edges). Each first-degree family network was analyzed with the prePRIMUS pipeline built into PRIMUS^92^ using the default settings to produce improved IBD estimates for the relationships within each family network and capture close relationships that span more than one ancestral superclass that were not captured in the ancestry-version IBD estimates. The two versions of IBD estimates were combined in the form of PLINK.genome file and subject to summary analysis after removing overly related samples with more than 100 close relatives (PI_HAT > 0.1875) or 25,000 relatives (PI_HAT > 0.08). All samples in the predicted first-and second-degree relationships were removed to generate the maximum unrelated data set for further analysis.

### Gene-and exon-level mutation rate calculations

Mutation rates were estimated using previously described methodology^93^. Rates for a particular trinucleotide context and methylation levels at CG sites were derived from gnomAD supplementary materials^2^. We assigned CpG methylation levels as gnomAD did with 3 tiers of methylation corresponding to <0.2, between 0.2-0.6, and >0.6 as low, medium, and high categories, respectively. Trimer contexts around every possible SNV substitution in coding regions were generated with bedtools v2.30.0 using coordinates from Ensembl GFF v100 and reference sequence GRCh38. To estimate pLOF mutation rates across a gene, trimer contexts were established for every pLOF SNV variant (stop-gained, splice acceptor, and splice donor) and matched to the gnomAD context-and methylation-specific rates. Transcript-and exon-level mutation rates were calculating by summing rates for all encompassed variants.

### Gene constraint (s_het_, heterozygous selection coefficient)

Mutations that cause deleterious effects with negative consequences on reproductive fitness are less likely to be transmitted to subsequent generations. These sites will experience fewer mutations than expected by chance and may be enriched in genes that perform biologically important functions. In a population genetics framework, selective pressure is modeled relative to fitness of homozygous carriers of the ancestral allele. Coefficients representing selection, *s*, and heterozygosity, *h*, of the variant’s impact on fitness, *w*, are modeled for heterozygous and homozygous carriers of the alternate allele as *w*=1*-hs* and 1-*s*, respectively. Cassa et al^23^ developed an approach to model a heterozygous selection coefficient for *hs*, i.e., s_het_, by treating this parameter as a random variable in a Bayesian joint-probability model alongside gene-specific SNV pLOF counts, *n*, for each gene. Allele counts for stop-gained, splice acceptor, and splice donor variants were summed across a gene following stringent filtering for variants that pass QC, LOFTEE high confidence, and mean depth ≥ 20 across the gene. The underlying data *n* is Poisson distributed with E[n] = (Nμ)/shet, where N is the number of chromosomes in the population supplying allele counts, and μ is a gene-specific mutation rate calculated as described above.

Here, we adapt the model from Cassa et al. and we jointly estimate distributions for the data, priors, and values of hyperparameters *a* and *b* in a hierarchical manner using pLOF variants with gene-wide cumulative MAF (cMAF) <0.001. As in Cassa’s original formulation, we set an inverse Gaussian prior on s_het_. Similarly, we separate the genes into 3 terciles based on mutation rates such that each tercile has different values of *a* and *b* tailored to low, medium, and high mutability. Hyperparameters *a* and *b* are modeled alongside s_het_ with inverse Gamma priors (shape=scale=1). We perform MCMC sampling with a Bayesian estimator, RStan^94^ using 30,000 iterations, discarding the initial 6,000 samplings, and thinning every 3 for 8,000 final iterations. Our application produces results on publicly available ExAC data that closely track the original Cassa et al. method in terms of accurately estimating hyperparameters and replicating their estimates, giving us confidence that our fully Bayesian formulation is appropriate.

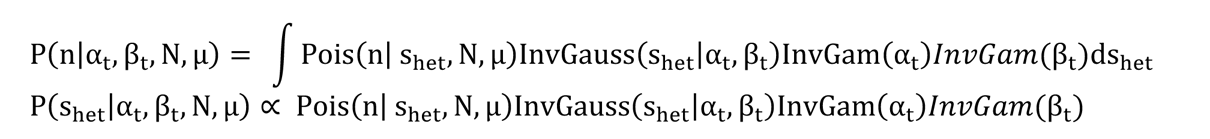

Population genetics theory suggests that, assuming strong selection and a large population, the frequency of rare, deleterious loss-of-function mutations can be estimated as *x = μ /(hs)*^24,95^. Weghorn et al.^24^ demonstrates that Bayesian estimates of s_het_ do not introduce additional variance compared with the deterministic mutation–selection balance approximation that underlies the Cassa et al. theoretical framework. The authors conclude that s_het_ is robust to the effects of genetic drift for most genes, except those under weak levels of selection starting near s_het_ around 0.02 and becoming more prominent at s_het_ ≤ 0.01. Therefore, our estimates of s_het_ are reasonable for our multi-ethnic, combined cohort analysis.

### Gene lists

Mouse knockout data from IMPC (Data Release 15.0, www.mousephenotype.org)^96^ and MGI were downloaded on 02/07/23; gene and term names were matched with human gene synonyms and Mouse Phenotyping Ontology (MPO) terms using downloadable lists available from MGI (http://www.informatics.jax.org/downloads/reports/index.html). Data from MGI were filtered to null/knockout phenotypes and term names including “lethality” were deemed lethal.

Similarly, data from IMPC were filtered for phenotypic effects with p<0.05 and lethal knockouts were determined using keywords “lethality” in the term name and “viability” in the procedure name. For both mouse KO datasets, the top level MPO terms MP:0010768 and MP:0005380 corresponding to mortality and embryo phenotypes were considered lethal. Cell essentiality screen associations were obtained from dbNSFP 4.3^97^ which in turn gathered data from 3 studies on CRISPR and large-scale mutagenesis assays^98–100^

Genes with variants in ClinVar^66^ (downloaded 0503/13/2023) labeled pathogenic or likely pathogenic, and in HGMD v2022.4 labeled disease mutations (DM) or disease-associated with additional supporting functional evidence (DFP) with high confidence were considered to have human disease associations. In addition, we used data from the DECIPHER developmental disease database (DDD) downloaded on 03/31/2022 and the COSMIC cancer gene census v95 (729 genes, tiers 1 and 2)^101^. Genes known to cause phenotypes with haploinsufficient, autosomal recessive, or autosomal dominant inheritance were downloaded from https://github.com/macarthur-lab/gene_lists and were originally collated from ClinGen or published literature.

### List of genes with homozygous pLOF variants

Genes with homozygous pLOF variants were culled from the Genome Aggregation Database (gnomAD) exomes (r2.1.1) and genomes (r3.1.2) data^2^, the NHLBI Trans-Omics for Precision Medicine program (TOPMED, Freeze 8)^19^ and the variant data included in the three publications, Pakistan Risk of Myocardial Infarction Study^35^, East London Genes and HEALTH^35^, and a whole genome study of Icelanders^37^. We annotated variant files from these different sources using our internal annotation pipeline and compiled a list of human genes with homozygous pLOF variants. Datasets not in HG38 human genome reference were transformed to HG38 coordinates using PICARD LiftoverVcf. Variants were further filtered with LOFTEE and MAF < 0.01 based on cohort-specific frequencies, while variants in RGC-ME were filtered to MAF<0.01 based on allele frequencies generated using 824k unrelated samples. For further consistency, we conducted our analyses on canonical transcripts and non-chromosome Y genes only. 4,874 genes have homozygous carriers of rare, alternate pLOF alleles across these 6 external datasets following these filters.

### MTR calculation

MTR is calculated within a sliding window with window sizes varies in 11 codons, 21 codons and 31 codons, moving by one codon each step. Smaller window sizes generate finer resolution of constrained regions using larger-scale data. Sliding windows in the beginning and end of the transcript is truncated until the window size is met. The expected variation is all possible variants in given transcripts annotated by VEP. The observed variation is tallied from 824K unrelated samples in RGC-ME data. The proportion of missense to the sum of missense and synonymous variants in the sliding window is calculated with both observed and expected data and MTR is the ratio of observed proportion to the expected as shown in the following formula. Correction for false discovery rate (FDR) was computed on all variants with MTR values and significant windows/variants passed an FDR<0.1 threshold. Percentile ranks were determined for all variants in canonical transcripts.

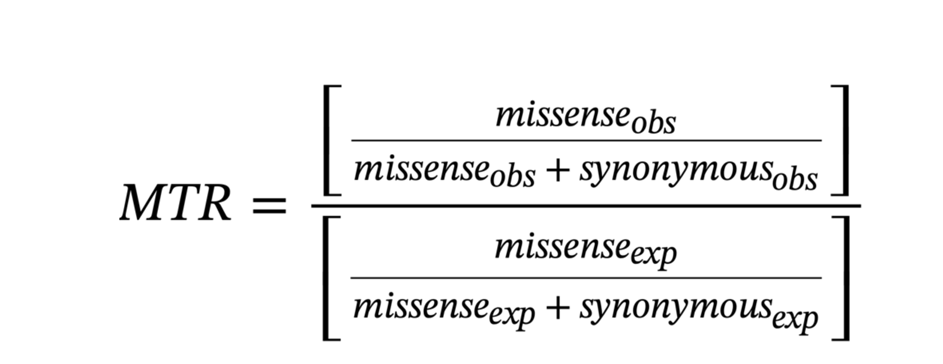

### MTR validation

MTR identified missense risk variants using the ClinVar variant database. Pathogenic and benign variants were extracted from ClinVar, which are identified as pathogenic/likely pathogenic and benign/likely benign and with at least “single submitter” as indicated by ClinVar^66^. Protein domain annotation was extracted from UniProt.

### Population differentiation

PLINK2 was used to calculate FST for each population pair at per-variant level. Each population was also compared to all other aggregated populations. Genome-wide summary statistics for all binary and quantitative traits for East Asian datasets were downloaded from Biobank Japan PheWeb (https://pheweb.jp/downloads). The data obtained from Biobank Japan was standardized and munged by mapping effect sizes to alternate alleles. Genome-wide summary statistics from UKB was used to identify association signals for differentiated variants in AFR, SAS and AMR.

### MAPS-derived threshold for splicing score

We adapted scripts from https://github.com/pjshort/dddMAPS to compute updated MAPS metrics, which adjusts for both mutation rate and methylation level in calculating the proportion of singletons. The mutation rate table was obtained from gnomAD^2^, with improved estimation of mutation rate by adjusting for methylation level compared with previous estimates^4^. For both SpliceAI and MMSplice, we set up a list of prediction score thresholds, ranging from 0.1 to 0.99, with step size of 0.2; For each threshold, we calculated the MAPS scores for both sets of variants that either pass or fail the threshold, separately. For each model, the threshold where the ‘passing’ variants can achieve a comparable deleteriousness as ‘most deleterious’ missense variants (predicted as deleterious variants by 5/5 deleterious prediction models) was chosen as the splicing score threshold for that model. Missense scores were derived from SIFT, Polyphen2_HDIV, Polyphen2_HVAR, LRT, and MutationTaster and obtained from dbNSFP v3.2. 5% of RGC-ME variants with SpliceAI score >0 and 3% of variants with MMSplice scores are identified as potentially deleterious variants that affect splicing (Fig. 5A, Fig. S8).

## Acknowledgments

Supported in part by the Intramural Research Program of the National Institute of Mental Health (ZIA-MH002843) and a grant from R01 NCI R01 CA157823. Ethical approval for the UK Biobank was previously obtained from the North West Centre for Research Ethics Committee (11/ NW/0382). The work described herein was approved by UK Biobank under application number 26041. Informed consent was obtained for all study participants. The authors thank everyone who made this work possible, the professionals from the member institutions who contributed to and supported this work, and most especially all research participants, without whom this research would not be possible. This study is funded by the Regeneron Genetics Center and Regeneron Pharmaceuticals

## Supplemental Tables

**Table S2:** s_het_ values for 16,704 genes and other annotations, including disease state (e.g. ClinVar or HGMD), LOEUF scores from gnomAD (2020 Supplementary Data), and coding sequence length. Values for computing s_het_ include variant count (n), total number of chromosomes (N_total), and mutation rate. S_het__lower and s_het__upper refer to the 2.5% and 97.5% of the posterior distribution, resulting in a 95% highest posterior density interval.

**Table S3:** Top results from gene set enrichment analysis for 1,241 genes deemed constrained using s_het_ cutoffs (mean>0.075, lower bound > 0.02). Gene set enrichment based on Reactome pathway genes performed using OxenrichR and filtered to adjusted p-value <0.05.

**Table S4:** List of genes with significant proportion of CDS in top 1, 5, 10, 15, and 20% percentile of exome wide MTR missense constraint scores (FDR<0.1) based on binomial test (π0=0.01, 0.05, 0.1, 0.15, and 0.2, respectively). Proportion of gene (count of amino acids with high MTR score, FDR<0.1, relative to total gene length) in each percentile compared with expected proportion based on corresponding null hypothesis. Resulting p-values are corrected for multiple testing with FDR (“BH”) and Bonferroni (“bonf”).

**Table S5:** List of 4,874 genes with rare (AAF<1%), homozygous pLOF variants. Value of homozygous and heterozygous carriers (“homAA”,“hetRA”), cumulative minor allele frequency (“cMAF”), and number of unique variants (“n”) are reported for the whole RGC-ME dataset including related individuals.

**Table S7.1:** List of highly differentiated variants (F_ST_ > 0.15) with MAF < 0.01 in Europeans.

**Table S7.2:** List of differentiated missense variants (F_ST_ > 0.05) with EUR MAF <0.01.

**Table S7.3:** Results of variant trait association analysis of AFR differentiated variants in UKB-AFR GWAS summary statistics.

**Table S7.4:** Results of variant trait association analysis of EAS differentiated variants in BBJ GWAS summary statistics.

## Notes

### Competing Interest Statement

K.S,X.., S.C, S.B,M.K, J.B, T.J, E.M, G.M, A.G, A.M, B.B, S.G, L.H, A.M, A.L, M.D.K, D.S., J.S., J.B., S.G, A.D, V.R, A.L, J.R.V, M.C, T.T, H.M.K, J.O, A.R.S, M.L.C, M.N, A.B, G.R.A, J.M., J.G.R, W.S, S.B are current employees and/or stockholders of Regeneron Genetics Center or Regeneron Pharmaceuticals.

### Summary of Updates

Added resource URL Updated Supplementary appendix

