## Supplementary Figures and tables for "A deep catalog of protein-coding variation in 985,830 individuals"

### Supplementary table 1: per-ancestry sample counts

| Description | >50% ancestry | Includes related individuals | Total samples | AFR | AMR | EAS | EUR | MEA | SAS | UNK | Non-EUR | Non-EUR % | Female % | Analysis use case |
|---|---|---|---|---|---|---|---|---|---|---|---|---|---|---|
| related analysis set | Y | Y | 985830 | 55497 | 85133 | 5884 | 724165 | 2249 | 30797 | 82105 | 179560 | 0.1821 | 0.5751 | Human knockout survey |
| unrelated analysis set | Y | N | 824159 | 49239 | 54563 | 5712 | 618161 | 2149 | 29521 | 64814 | 141184 | 0.1713 | 0.5644 | Variant survey (incl. pathogenic variants and ACMG genes), constraint metrics ($s_{het}$ and MTR), MAPS, splice scores, $F_{ST}$ |
| unrelated proportional set | N | N | 824159 | 49619.53 | 54628.88 | 15042.03 | 633466.94 | 3945.75 | 40723.87 | 26732 | 163960 | 0.1989 | | Proportional sample sizes for browser, fine-scale ancestry AF, and Figure 1 |

**Supplementary Table 1.** Sample subsets of RGCME used in different analyses, depending on whether related samples were appropriate or not, and if ancestry assignment was proportional or assigned based on maximum-likelihood probability above 50%. Total number of samples in analysis set and divided by ancestry are shown.

### Per individual variant counts: Supplementary Figure 1

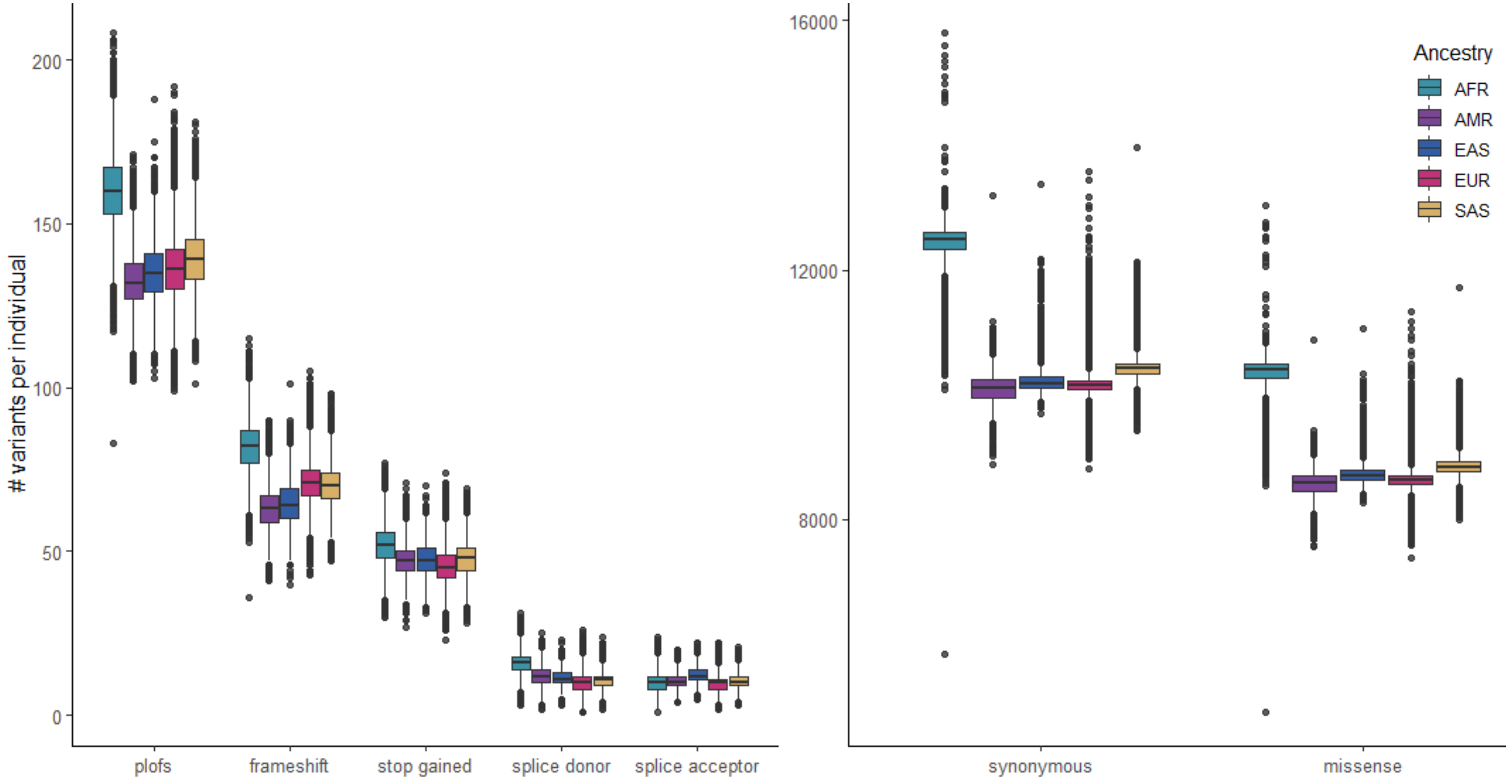


**Supplementary Figure 1.** Box plot of variant counts observed in each unrelated sample in RGCME. Horizontal lines in the box represent the 25%, 50%, and 75% percentile of the distribution of counts and points represent outliers. Per-individual counts are divided by ancestral assignment (>50% probability).

### Mutation saturation: Supplementary Figure 2

**Supplementary Figure 2**: Mutation saturation survey in RGC-ME data

Counts are based on variant-transcript pairs.

Methylation level*: mean methylation values across tissues of >0.65, 0.2-0.65, <0.2 correspond to methylation level of 2, 1, 0 respectively

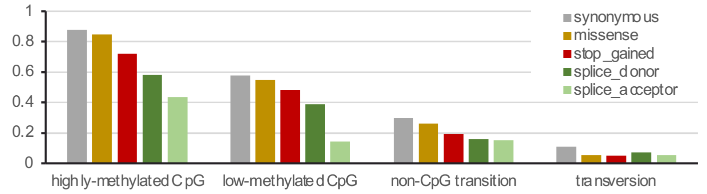


| Effect | Variant type | Methylatio level* | # Observed | # Singletons | # Possible | % Observed | % Singletons |
|---|---|---|---|---|---|---|---|
| **synonymous** | CpG | 0 | 92,633 | 21,678 | 170,780 | 54.24% | 23.40% |
| | | 1 | 38,104 | 4,402 | 55,738 | 68.36% | 11.55% |
| | | 2 | 406,216 | 5,877 | 463,866 | 87.57% | 1.45% |
| | non-CpG | | 3,153,425 | 1,080,198 | 10,524,791 | 29.96% | 34.25% |
| | transversion | | 1,248,227 | 473,981 | 11,354,674 | 10.99% | 37.97% |
| **missense** | CpG | 0 | 136,027 | 35,140 | 268,643 | 50.63% | 25.83% |
| | | 1 | 62,573 | 8,329 | 94,795 | 66.01% | 13.31% |
| | | 2 | 826,776 | 26,766 | 977,530 | 84.58% | 3.24% |
| | non-CpG | | 5,183,884 | 1,889,827 | 19,919,347 | 26.02% | 36.46% |
| | transversion | | 4,949,847 | 2,037,927 | 52,377,079 | 9.45% | 41.17% |
| **splice_acceptor** | CpG | 0 | 1 | 0 | 6 | 16.67% | 0.00% |
| | | 1 | | | 1 | 0.00% | |
| | | 2 | 16 | 0 | 37 | 43.24% | 0.00% |
| | non-CpG | | 56,418 | 25,940 | 370,116 | 15.24% | 45.98% |
| | transversion | | 40,732 | 19,909 | 740,322 | 5.50% | 48.88% |
| **splice_donor** | CpG | 0 | 171 | 63 | 577 | 29.64% | 36.84% |
| | | 1 | 176 | 28 | 318 | 55.35% | 15.91% |
| | | 2 | 2,348 | 146 | 4,028 | 58.29% | 6.22% |
| | non-CpG | | 58,917 | 25,549 | 365,238 | 16.13% | 43.36% |
| | transversion | | 51,537 | 24,715 | 740,321 | 6.96% | 47.96% |
| **stop_gained** | CpG | 0 | 1,582 | 553 | 4,237 | 37.34% | 34.96% |
| | | 1 | 1,906 | 286 | 3,017 | 63.18% | 15.01% |
| | | 2 | 42,460 | 4,356 | 58,817 | 72.19% | 10.26% |
| | non-CpG | | 155,714 | 71,309 | 808,575 | 19.26% | 45.79% |
| | transversion | | 162,186 | 77,432 | 3,294,877 | 4.92% | 47.74% |
| **frameshift** | insertion | | 427,053 | 221,112 | | | 51.78% |
| | deletion | | 185,217 | 94,613 | | | 51.08% |

### Supplementary Figure 3

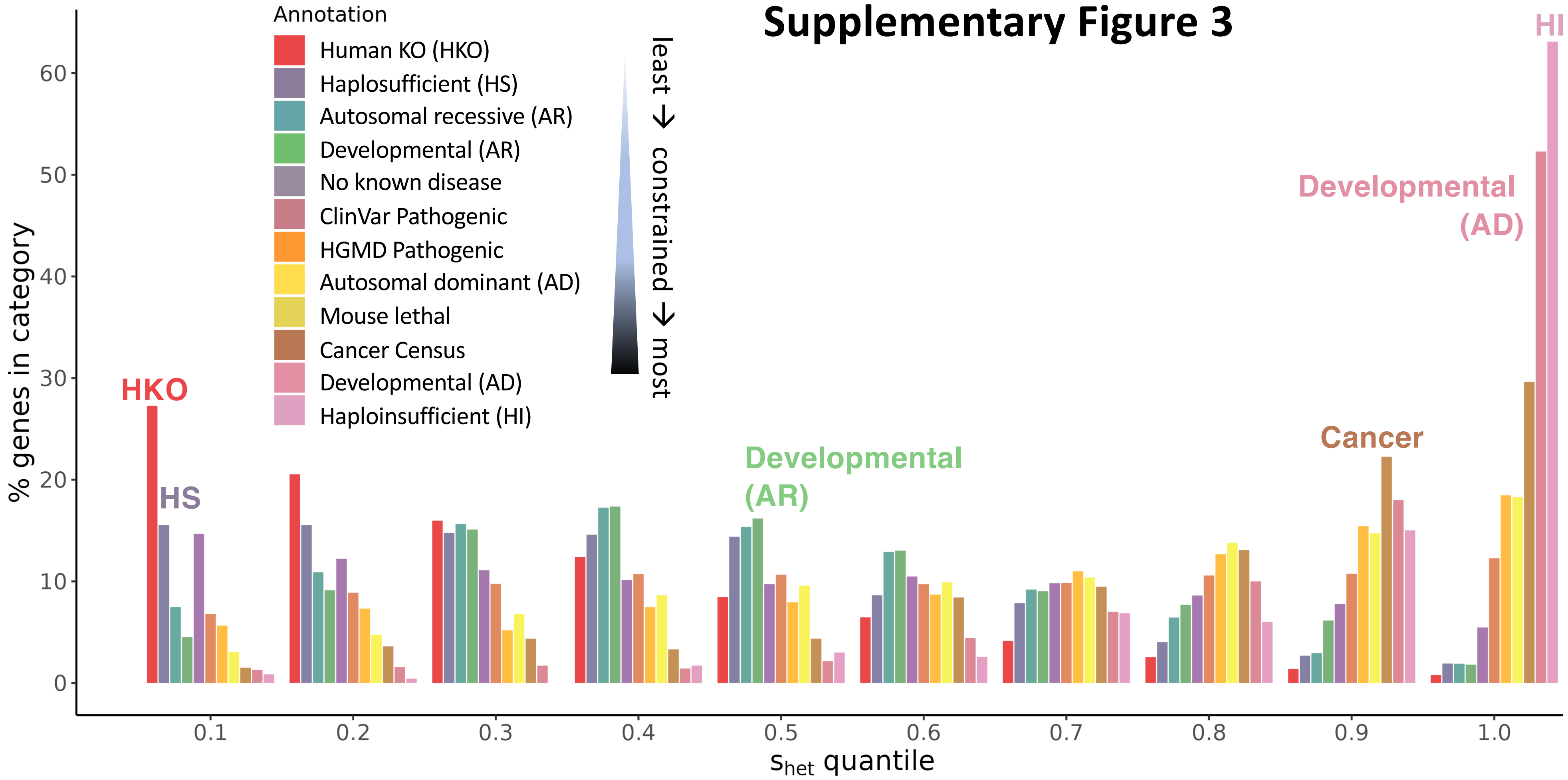


**Supplemental Figure 3**: Proportion of genes in each disease/essentiality annotation list (gathered from published literature and databases) that correspond to each $s_{het}$ decile.

### Supplementary Figure 4

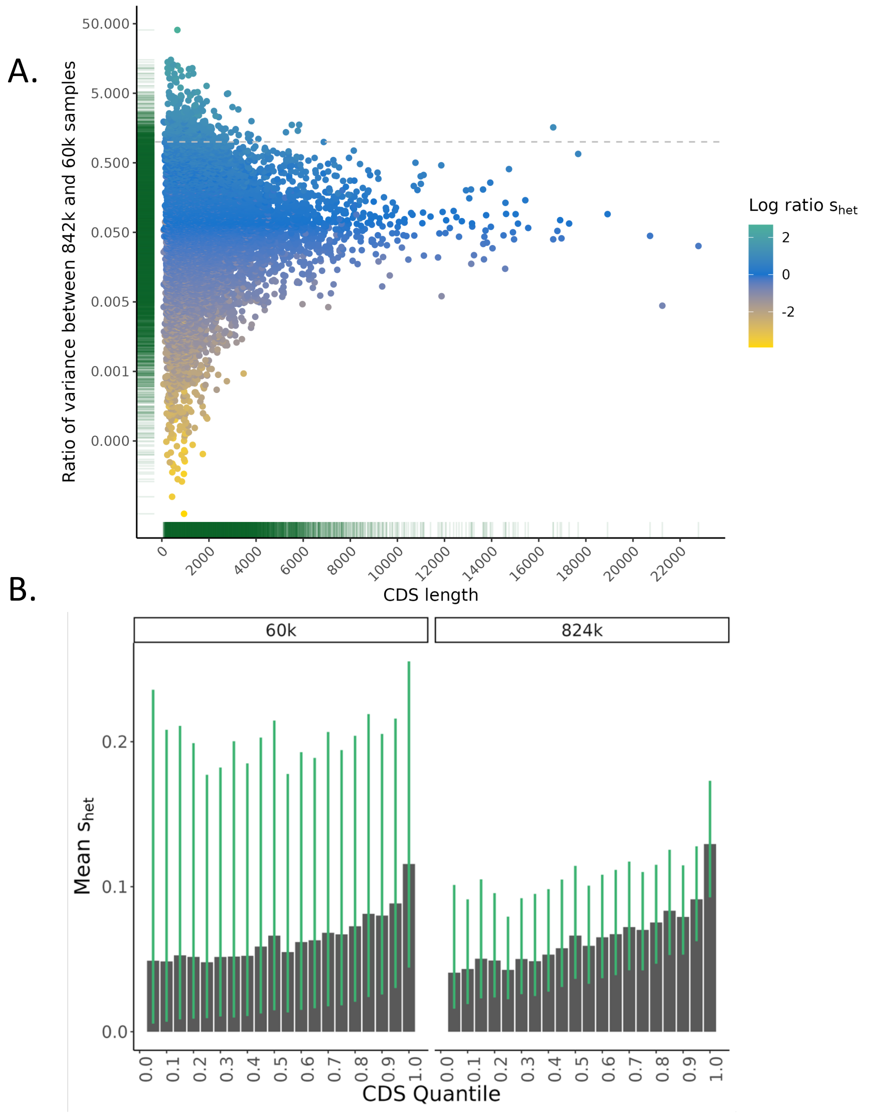


**Supplemental Figure 4**: A. Ratio of variances for $s_{het}$ calculated on full dataset (824k) and randomly downsampled set of 60,000 individuals. The mean ratio around 0.05 suggests that gene-level variance from the full dataset is 20x smaller than variance for the same gene using the downsampled set. This is the case despite similar, or even higher, $s_{het}$ mean estimates as shown by the color bar. B. Mean and 95% HPD (green bar) $s_{het}$ estimates for genes in each CDS length quartile for downsampled 60k (left) and full 824k (right) samples.

### Supplementary Figure 5

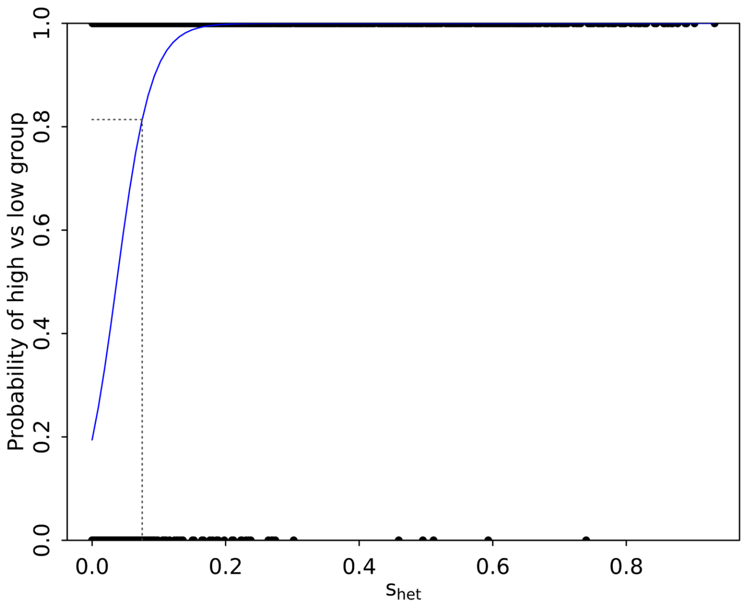


**Supplementary Figure 5**. Logistic regression for probability that gene with $s_{het}$=*x* will be in group=1, "highly constrained" represented by haploinsufficient, developmental disorder autosomal dominant, and mouse lethal genes or group=0 "not highly constrained" represented by haplosufficient and genes with rare, homozygous pLOFs in RGC-ME. Dotted lines for genes with $s_{het}$ = 0.075 resulting in probability of 81% and of being in the high group (with precision=0.95)

### Supplementary Figure 6

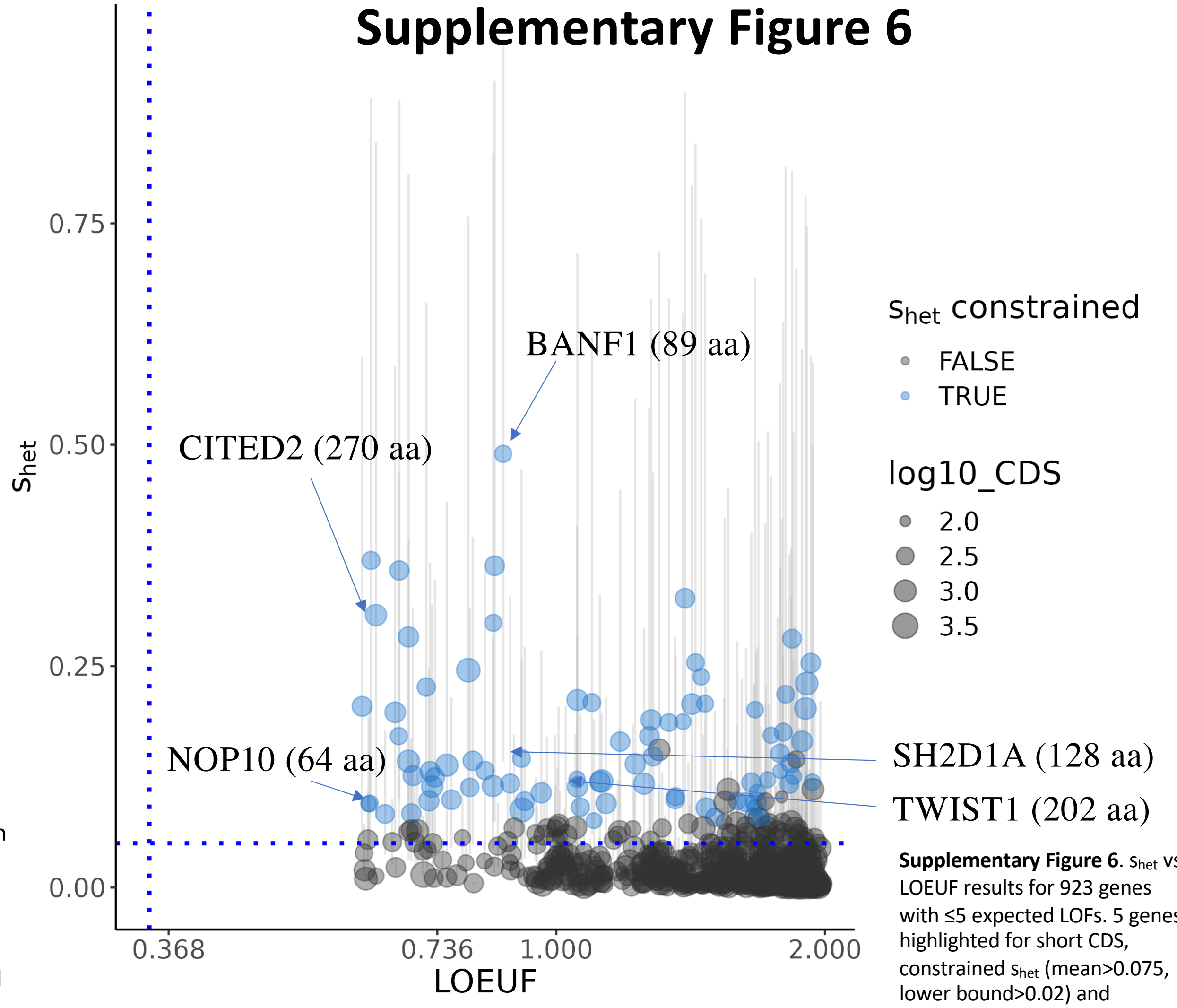


**Supplementary Figure 6**. $s_{het}$ vs LOEUF results for 923 genes with ≤5 expected LOFs. 5 genes highlighted for short CDS, constrained $s_{het}$ (mean>0.075, lower bound>0.02) and unconstrained LOEUF.

### Supplementary Figure 7

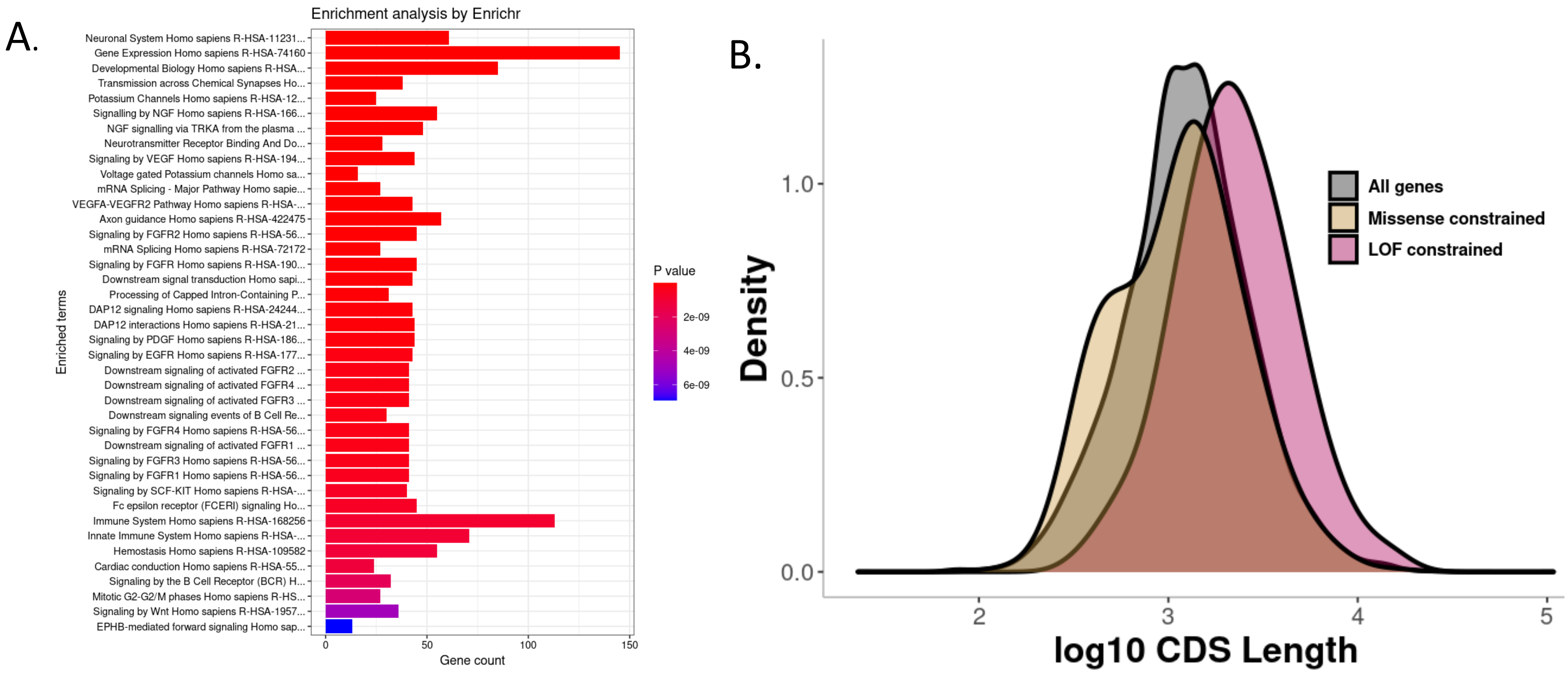


**Supplementary Figure 7:**A. Reactome pathway enrichment of 1,818 genes with human-derived most constrained missense sites that are not conserved across species. C. The CDS length comparison of missense constraint genes and LoF constraint genes. Missense constrained genes include 292 genes with $s_{het}$ score <0.075 and 165 genes without $s_{het}$ estimations. LOF constrained genes are 641 genes with $s_{het}$ score >=0.075 and insignificant enrichment of top 1% MTR variants (FDR>0.1) or null MTR value. All genes are in total of 19,644 genes with canonical transcripts shown in the grey curve. The analysis is based on canonical transcripts.

**Table S6 – ancestral breakdown**

On left, number of individuals per ancestry in HKO analysis, including related individuals, based on continental ancestry with probability assignment >50%. On right, the number of genes with greater than 1, 5, and 10 carriers observed with rare, homozygous alternate allele variants in LOFs.

| | | # of genes with | | |
|---|---|---|---|---|
| | Total samples | ≥1 carrier | ≥5 carriers | ≥10 carriers |
| AFR | 55,497 | 1,098 | 403 | 298 |
| AMR | 85,133 | 1,287 | 160 | 83 |
| EAS | 5,884 | 318 | 57 | 35 |
| EUR | 724,165 | 2,590 | 698 | 409 |
| SAS | 30,797 | 2,102 | 255 | 108 |
| ALL | 985,830 | 4,874 | 1,457 | 915 |

### Supplementary Figure 8

| Gene | Closest paralog | % target match | # paralogs |
|---|---|---|---|
| CYP2C9 | CYP2C19 | 91.4 | 16 |
| CYP2D6 | CYP2D7 | 94.0 | 16 |
| CYP1A1 | CYP1A2 | 72.3 | 2 |
| CYP2E1 | CYP2C18 | 56.2 | 16 |
| CYP4F22 | CYP4F3 | 64.0 | 12 |
| CYP4F2 | CYP4F3 | 87.7 | 12 |
| CYP2B6 | CYP2A13 | 53.6 | 16 |
| CYP2A7 | CYP2A6 | 93.7 | 16 |
| CYP2S1 | CYP2A13 | 48.4 | 16 |

| | Cumulative pLOF AAF | | | |
|---|---|---|---|---|
| **Gene** | **AFR** | **AMR** | **EUR** | **SAS** |
| **CYP2C19** | 8.56E-04 | 2.98E-03 | 2.97E-04 | 3.35E-03 |
| **CYP2C9** | 1.16E-02 | 8.34E-04 | 3.06E-04 | 5.58E-04 |
| **CYP2D6** | 3.79E-04 | 1.35E-04 | 1.32E-04 | 6.12E-04 |
| **CYP2A6** | 2.04E-02 | 2.82E-04 | 2.29E-04 | 1.76E-04 |
| **CYP1A2** | 2.61E-04 | 3.88E-04 | 5.53E-04 | 1.14E-03 |

**Supplemental Figure S6:** A closer look at the CYP family of genes, which were highly represented among putative gene KOs. A: The number of paralogous genes among CYP species identified with HKOs and the % target match of the closest paralog. B. Cumulative allele frequencies for pLOF variants observed in CYP genes with HKOs. AAF computed using RGCME samples of continental ancestries (>50% probability in related samples) for these genes demonstrate differences in AAF across population groups.

### Supplementary Figure 9.1

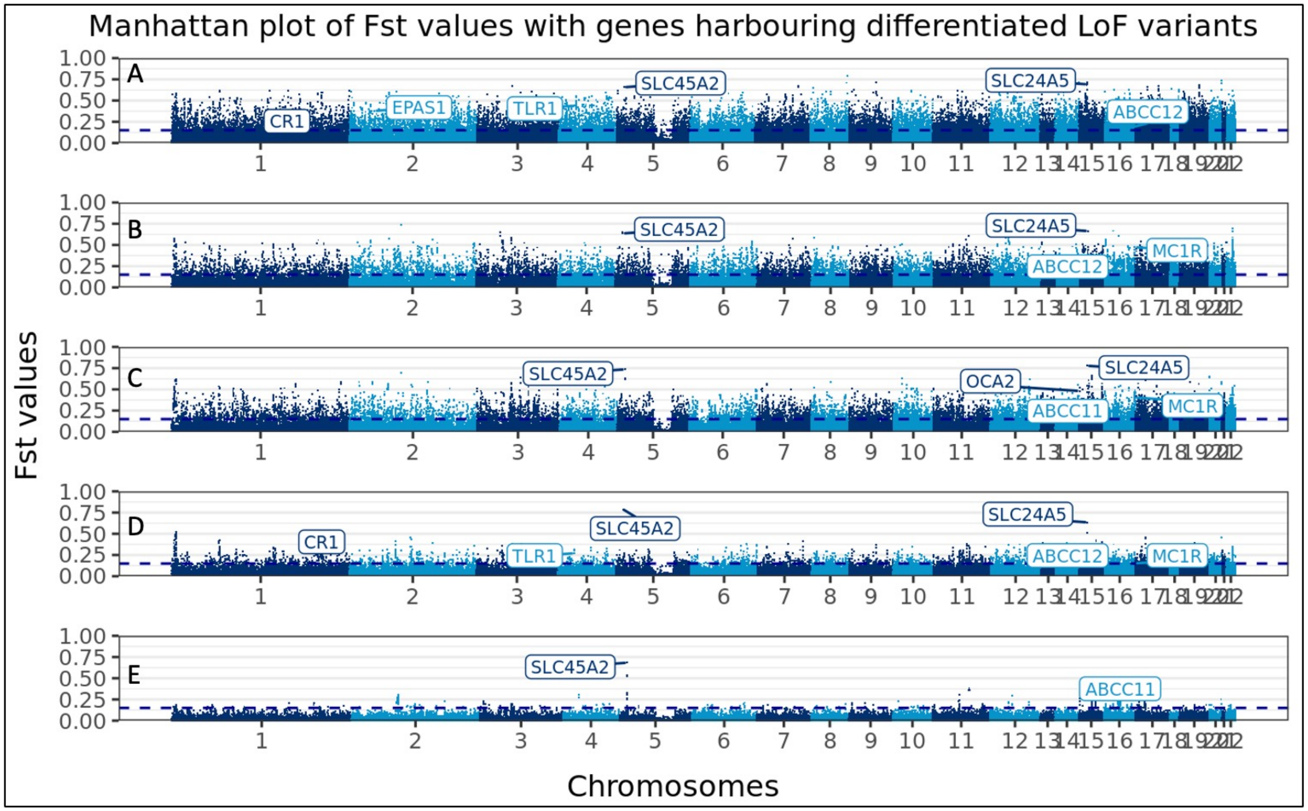


**Supplementary Figure 9.1**. Manhattan plots of differentiated variants in (A) AFR, (B) AMR, (C) EAS, (D) EUR, and (E) SAS. Dotted blue line depicts Fst > 0.15. Annotated genes are known to be under selection. Many variants were found to be highly differentiated in AFR, EAS, and AMR, in comparison to SAS.

### Supplementary figure 9.2

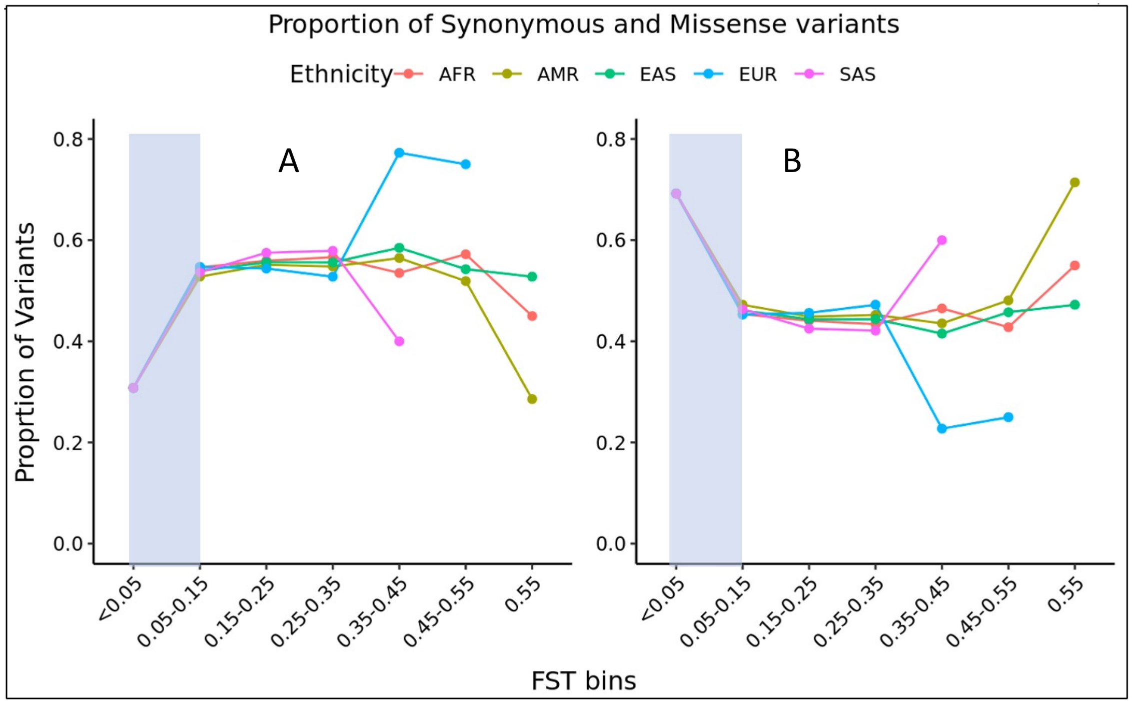


Fig 9.2: Distribution of (A) Synonymous variants and (B) and missense variants by different $F_{ST}$ bins. Proportion of missense variants drops (blue shaded area) in comparison to synonymous variants for the differentiated variants in higher $F_{ST}$ bins. This distribution is similar among different populations. For very high $F_{ST}$ bins the distributions among different populations because of variability due to smaller number of variants

### Supplementary Figure 10

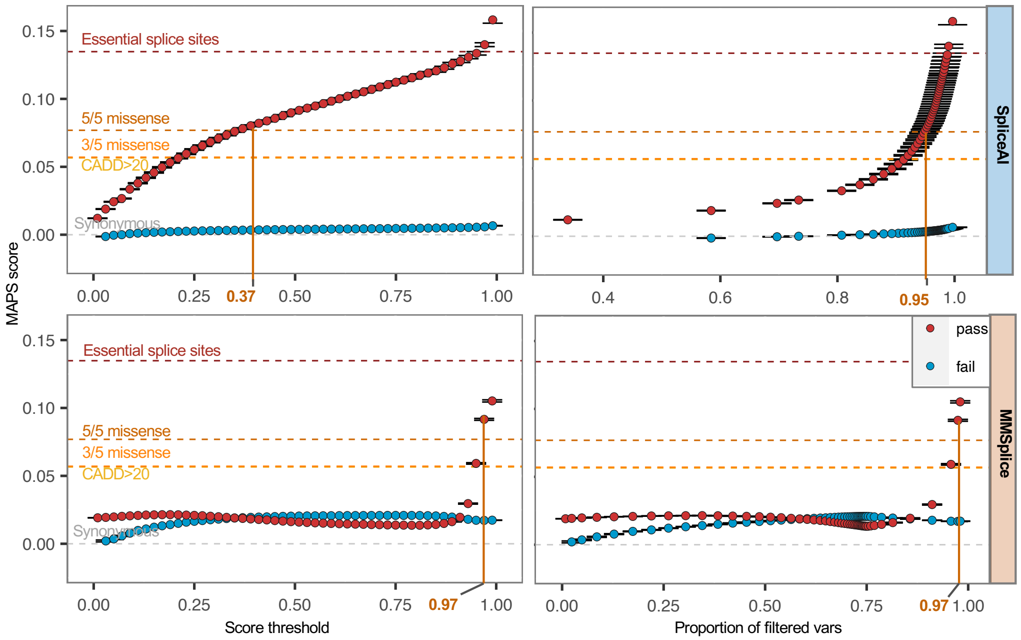


**Supplementary Figure 10.** Schematic of MAPS-derived filters for SpliceAI and MMSplice. To systematically evaluate the deleteriousness of cryptic splice variants identified by each threshold, a list of prediction score thresholds were set up for both spliceAI and MMSplice, ranging from 0.1 to 0.99, with step size of 0.2. For each threshold, we divided variants into two sets: the set of variants passed the threshold, represented as red dots, and the set of variants that failed the filter, represented as blue dots. The y-axis represents MAPS score for each set of variants; the X-axis shows the tested score threshold (left panels) or the percent of variants have been filtered under each threshold (right panels). All MAPS scores were calculated based on the QC passed variants with splicing prediction scores in RGC-ME unrelated samples.

### Supplementary Figure 11

A.

| Category | Total | SpliceAI (score>0.35) Counts | SpliceAI (score>0.35) % | MMSplice (score>0.97) Counts | MMSplice (score>0.97) % | Intersection set Counts | Intersection set % |
|---|---|---|---|---|---|---|---|
| HC.SS | 216,702 | 189,749 | **87.56** | 175,122 | **80.81** | 163,551 | **75.47** |
| LC.SS | 8,425 | 4,862 | **57.71** | 5,092 | **60.44** | 4,225 | **50.15** |
| splice_region | 1,117,939 | 131,703 | 11.78 | 198,699 | 17.77 | 90,884 | 8.13 |
| stop_gained | 344,093 | 14,687 | 4.27 | 4,603 | 1.34 | 2,117 | 0.62 |
| start_lost | 30,525 | 276 | 0.90 | 249 | 0.82 | 45 | 0.15 |
| stop_lost | 12,466 | 84 | 0.67 | 82 | 0.66 | 3 | 0.02 |
| 5/5 missense | 2,953,321 | 42,182 | 1.43 | 27,841 | 0.94 | 12,112 | 0.41 |
| Other missense | 8,039,579 | 99,553 | 1.24 | 53,862 | 0.67 | 20,361 | 0.25 |
| synonymous | 4,699,177 | 40,738 | 0.87 | 3,674 | 0.08 | 34 | 7.24E-04 |
| Total | 17,422,227 | 523,834 | 3.01 | 469,224 | 2.69 | 293,332 | 1.68 |

B.

C.

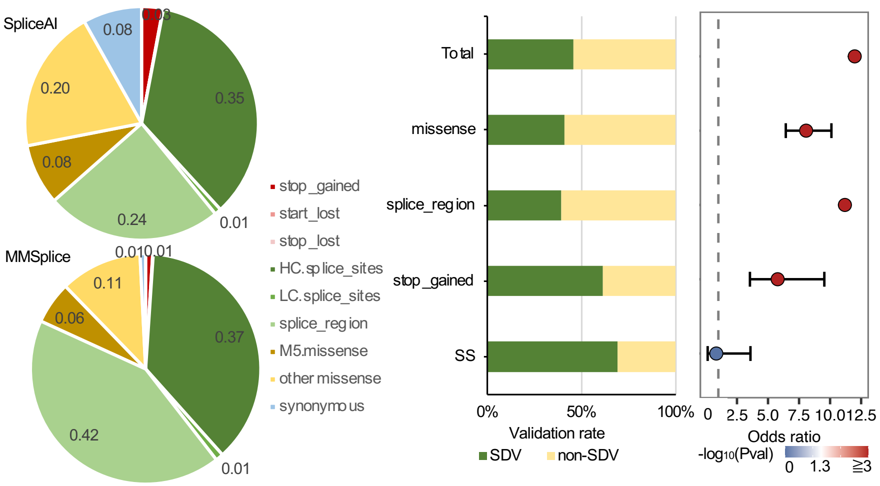


**Supplementary Figure 11**.

A. Summary of splice-disrupting coding variants in different functional categories using MAPS-defined prediction score thresholds for SpliceAI and MMSPlice.

B. Distribution of different functional categories of splice-disrupting coding variants. **HC:** High confidence **LC:** low confidence. These annotation tags are derived from LOFTEE

A. Empirical validation of MAPS scores.

- Left panel: Fraction of predicted splice-affecting variants (union set) that validated as splice disrupting variants (SDVs) by any of the three splice reporter assays
- Right panel: enrichment of predicted splice-affecting variants in SDVs compared to non-SDVs

**Table S8**: Summary of splice disrupting variants collected from three splicing reporter assays

| Assay | Total | SDVs |
|---|---|---|
| Vex-seq<br>Adamson et al. Genome Biology, 2018 | 1,960 | 796 (ΔPSI>0.05) |
| MaPSy<br>Soemedi et al. Nat Genetics, 2017 | 5,179 | 962 (FC>1.5, FDR<0.05) |
| MFASS<br>Cheung et al. Mol Cell, 2019 | 28,972 | 1,050 (almost complete loss of exon recognition) |
| Total | 36,067 | 2,806 |

### Supplementary Figure 12

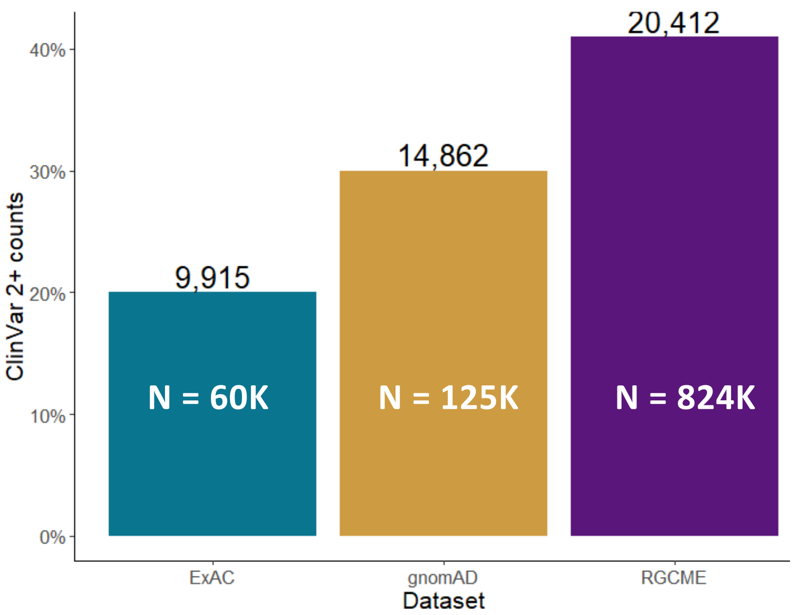


Supplementary Figure 12: Counts of pathogenic variants in ClinVar with high confidence (STAR≥2, total=50,036) observed in large-scale exome sequencing studies, including RGCME, gnomAD, and ExAC is indicted on the top of each bar. The number of individuals in each dataset is indicated inside each bar

### Table S9

| | | All variants |
|---|---|---|
| Known pathogenic (P) | **Genes** | 72 |
| | **Variants** | 2,829 |
| | – TTR and HFE | 2,812 |
| | **Reportable carriers** | 22,846 |
| | – TTR and HFE | 17,278 |
| | **Carrier rate** | 2.77% |
| | – TTR and HFE | 2.10% |
| *Likely pathogenic (LP) | **Genes** | 40 |
| | **Variants** | 1,407 |
| | **Reportable carriers** | 2,357 |
| | **Carrier rate** | 0.29% |
| **Total carrier rate** | | 3.06% |
| – TTR and HFE | | 2.38% |

Supplementary Table 9: Count of individuals and percent of total RGC-ME which comprise carriers of variants in ACMG reportable genes, excluding or including TTR and HFE. Carriers of LP variants have observed pLOFs in the canonical transcript of relevant genes.

### Supplementary Figure 13

Supplementary Figure 13: Counts of ClinVar variants with differential allele frequency between ancestral groups compared with EUR, i.e., fold-change = $AAF_{AFR}$ / $AAF_{EUR}$. Variants are classified in fold-change bins ranging from only observed in non-EUR, to elevated in non-EUR (fold change bins >1x), to only observed in EUR. Panels A-B depict VUS and C-D depict pathogenic variants. Panels B and D are zoom-in views of the AAF observed in non-EUR groups for variants that are >100x less frequent in EUR. For example, per sub-sample, ~500 variants have AAF={0.01,0.05} in AFR and 100x fold lower AAF in EUR.

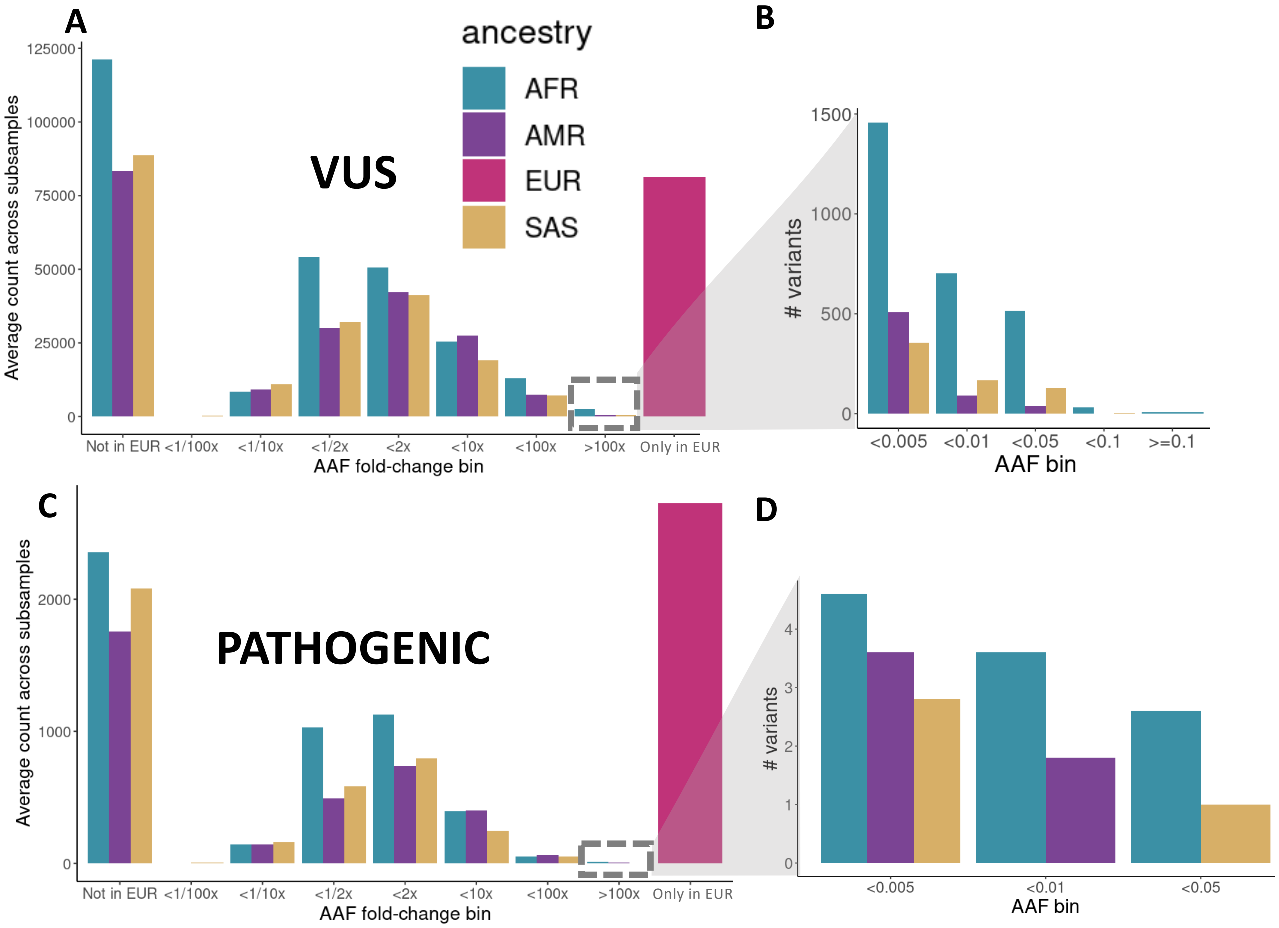

### Table S10

| | ClinVar VUS variants | | Observed missense in 824k | |
|---|---|---|---|---|
| MTR cutoff | Variant count | % of VUS missense | Variant count | % of observed missense |
| 1 percentile | 6,117 | 2.99% | 19,231 | 0.18% |
| 5 percentile | 24,511 | 11.98% | 196,236 | 1.88% |
| 10 percentile | 32,617 | 15.94% | 355,582 | 3.40% |
| 20 percentile | 68,432 | 33.44% | 823,726 | 7.87% |

Supplementary Table 10. The distribution of ClinVar VUS variants and missense variants observed in 824K unrelated samples in top MTR constrained regions (FDR<0.1).

### Supplementary Figure 14

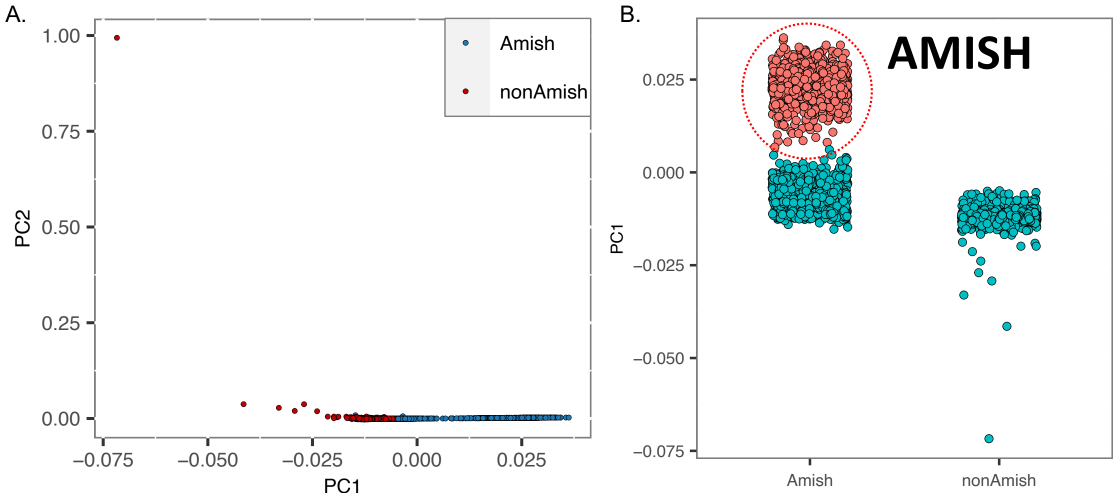


**Supplementary Fig 14:** A. Plots of the first two principal components derived from 1963 UKB and 1963 samples from predominantly Amish cohorts B. k-means clustering on PC1 to unambiguously identify the subset of Amish individuals (red cluster) who are distinct from the UKB European cluster (blue cluster)

### Supplemental Tables

**Table S2:** $s_{het}$ values for 16,704 genes and other annotations, including disease state (e.g. ClinVar or HGMD), LOEUF scores from gnomAD (2020 Supplementary Data), and coding sequence length. Values for computing $s_{het}$ include variant count (n), total number of chromosomes (N_total), and mutation rate. $S_{het}$_lower and $s_{het}$_upper refer to the 2.5% and 97.5% of the posterior distribution, resulting in a 95% highest posterior density interval.

**Table S3:** Top results from gene set enrichment analysis for 1,241 genes deemed constrained using $s_{het}$ cutoffs (mean>0.075, lower bound > 0.02). Gene set enrichment based on Reactome pathway genes performed using OxenrichR and filtered to adjusted p-value <0.05.

**Table S4:** List of genes with significant proportion of CDS in top 1, 5, 10, 15, and 20% percentile of exome wide MTR missense constraint scores (FDR<0.1) based on binomial test (π0=0.01, 0.05, 0.1, 0.15, and 0.2, respectively). Proportion of gene (count of amino acids with high MTR score, FDR<0.1, relative to total gene length) in each percentile compared with expected proportion based on corresponding null hypothesis. Resulting p-values are corrected for multiple testing with FDR ("BH") and Bonferroni ("bonf").

**Table S5:** List of 4,874 genes with rare (AAF<1%), homozygous pLOF variants. Value of homozygous and heterozygous carriers ("homAA","hetRA"), global allele frequency ("gMAF"), and number of unique variants ("n") are reported for the whole RGC-ME dataset including related individuals.

**Table S7.1:** List of highly differentiated variants ($F_{ST}$ > 0.15) with MAF < 0.01 in Europeans.

**Table S7.2:** List of differentiated missense variants ($F_{ST}$ > 0.05) with EUR MAF <0.01.

**Table S7.3:** Results of variant trait association analysis of AFR differentiated variants in UKB-AFR GWAS summary statistics.

**Table S7.4:** Results of variant trait association analysis of EAS differentiated variants in BBJ GWAS summary statistics.
