## Supplementary Appendix for "A deep catalog of protein-coding variation in 985,830 individuals"

### Regeneron Genetics Center


**Analytical Genetics and Data Science**
Gonçalo Abecasis, Manuel Allen Revez Ferreira, Joshua Backman, Kathy Burch, Adrian Campos, Lei Chen, Sam Choi, Amy Damask, Lee Dobbyn, Liron Ganel, Sheila Gaynor, Benjamin Geraght, Akropravo Ghosh, Christopher Gillies, Lauren Gurski, Joseph Herman, Eric Jorgenson, Tyler Joseph, Michael Kessler, Jack Kosmicki, Nan Lin, Adam Locke, Jonathan Marchini, Anthony Marcketta, Joelle Mbatchou, Arden Moscati, Priyanka Nakka, Aditeya Pandey, Anita Pandit, Charles Paulding, Jonathan Ross, Carlo Sidore, Eli Stahl, Maria Suciu, Timothy Thornton, Peter VandeHaar, Sailaja Vedantam, Scott Vrieze, Rujin Wang, Kuan-Han Wu, Bin Ye, Blair Zhang, Andrey Ziyatdinov, Yuxin Zou

**Clinical Informatics**
Amelia Averitt, Nilanjana Banerjee, Michael Cantor, Dadong Li, Sameer Malhotra, Justin Mower, Mudasar Sarwar, Deepika Sharma, Jeffrey C Staples, Jay Sundaram, Sean Yu, Aaron Zhang

**Genome Informatics & Data Engineering**
Xiaodong Bai, Suganthi Balasubramanian, Suying Bao, Boris Boutkov, Andrew Bunyea, Janice Clauer, Evan Edelstein, Gisu Eom, Sujit Gokhale, Alexander Gorovits, Ju Guan, Lukas Habegger, Alicia Hawes, Olga Krasheninina, Rouel Lanche, Vrushali Mahajan, Koteswararao Makkena, Adam J Mansfield, Evan K Maxwell, George Mitra, Mona Nafde, Sean O'Keeffe, Razvan Panea, Krishna Pawan Punuru, Tommy Polanco, Ayesha Rasool, Jeffrey G Reid, William J Salerno, Sanjay Sreeram, Benjamin Sultan, Kathie Sun, Lance Zhang

**Research Program Management & Strategic Initiatives**
Esteban Chen, Jaimee Hernandez, Marcus B Jones, Michelle G LeBlanc, Jason Mighty, Nirupama Nishtala, Nadia Rana, Jennifer Rico-Varela

**RGC Mangement & Leadership Team**
Gonçalo Abecasis, Aris Baras, Michael Cantor, Giovanni Coppola, Andrew Deubler, Aris Economides, Katia Karalis, Luca A Lotta, Lyndon J Mitnaul, John D Overton, Jeffrey G Reid, Alan Shuldiner, Katherine Siminovitch

**Sequencing & Lab Operations**
Christina Beechert, Erin D Brian, Laura M Cremona, Hang Du, Caitlin Forsythe, Zhenhua Gu, Kristy Guevara, Michael Lattari, Alexander Lopez, Kia Manoochehri, John D Overton, Manasi Pradhan, Raymond Reynoso, Ricardo Schiavo, Maria Sotiropoulos Padilla, Chenggu Wang, Sarah E Wolf

**Therapeutic Area Genetics**
Parsa Akbari, Anna Alkelai, Silvia Alvarez, Ariane Ayer, Antoine Baldassari, Jonas Bovijn, Jessie Brown, Giovanni Coppola, Tanima De, Peter Dombos, Adolfo Ferrando, Jan Freudenberg, Sahar Gelfman, Arthur Gilly, Sujit Gokhale, Sarah Graham, Aysegul Guvenek, Mary Haas, Jin He, George Hindy, Brian Hobbs, Amit Joshi, Manav Kapoor, Hossein Khiabanian, Vijay Kumar, Luca A Lotta, Priyanka Nakka, Jacqueline Otto, Billy Palmer, Neel Parikshak, Veera Rajagopal, Moeen Riaz, Juan Rodriguez-Flores, Alan Shuldiner, Katherine Siminovitch, Olukayode Sosina, Kayode Sosina, Luanluan Sun, Gannie Tzoneva, Niek Verweij, Cristen J Willer, Bin Ye

### RGC-ME Cohort Partners

**Accelerated Cures**
David Gwynne
**Albert Einstein College of Medicine**
Nir Barzilai, Yousin Suh, Zhengdong Zhang
**Case Western Reserve University**
Jonathan L. Haines
**Center for Non-Communicable Diseases, Karachi, Pakistan**
Danish Saleheen
**Cincinnati Children's Hospital**
Ken Kaufman, Leah Kottyan, Lisa Martin, Marc Rothenberg
**Columbia University**
Abdullah Ali, Azra Raza
**Duke University**
William E. Kraus, Christopher B. Newgard, Svati Shah
**Flinders University of South Australia**
Jamie Craig, Alex Hewitt
**Florey Institute of Neuroscience and Mental Health, on behalf of the Australia and New Zealand MS Genetics Consortium**
Trevor J. Kilpatrick, Justin Rubio
**Geisinger Heath System**
Lance J. Adams, Jackie Blank, Dale Bodian, Derek Boris, Adam Buchanan, David J. Carey, Ryan D. Colonie, F. Daniel Davis, Dustin N. Hartzel, Melissa Kelly, H. Lester Kirchner, Joseph B. Leader, David H. Ledbetter, J. Neil Manus, Christa L. Martin, Raghu P. Metpally, Michelle Meyer, Tooraj Mirshahi, Matthew Oetjens, Thomas Nate Person, Christopher Still, Natasha Strande, Amy Sturm, Jen Wagner, Marc Williams
**Griffith University, on behalf of the Australia and New Zealand MS Genetics Consortium**
Simon Broadley
**Indiana University School of Medicine**
Naga Chalasani, Tatiana Foroud, Suthat Liangpunsakul, Lois B. Travis
**Kaiser Permanente**
Lori Sakado, John Witte
**Loyola University**
Heather Wheeler
**Lundquist Institute**
Jerome I. Rotter, Robert Weinreb
**Mayo Clinic**
Kostantinos Lazaridis
**Monash University, Melbourne, VIC, Australia, on behalf of the Australia and New Zealand MS Genetics Consortium**
Vilija Jokubaitis
**Mount Sinai School of Medicine**
Erwin Bottinger, Judy Cho

**Murdoch University, on behalf of the Australia and New Zealand MS Genetics Consortium**
Marzena J. Fabis-Pedrini, Allan G Kermode
**National Autonomous University of Mexico**
Jesus Alegre-Diaz, Pablo Kuri-Morales, Roberto Tapia-Conyer
**National Institute of Mental Health, NIH**
Francis McMahon
**Northwestern University**
Adam Gordon, Maureen Smith, John Varga
**Perron Institute for Neurological and Translational Science, Perth, Australia, on behalf of the Australia and New Zealand MS Genetics Consortium**
Allan G Kermode
**The University of New Castle, Australia, on behalf of the Australia and New Zealand MS Genetics Consortium**
Jeanette Lechner-Scott, Rodney J. Scott
**University of Alabama at Birmingham**
S. Louis Bridges, Robert Kimberly
**University of California, Los Angeles**
Marlena Fejzo
**University of Chicago**
Eileen Dolan, Omar El-Charif
**University of Colorado School of Medicine**
Richard A. Spritz
**University of Maryland School of Medicine**
Elliot Hong, Braxton Mitchell
**University of Melbourne, Victoria, Australia, on behalf of the Australia and New Zealand MS Genetics Consortium**
Stephen Leslie
**University of Miami**
Margaret A. Pericak-Vance
**University of Michigan Medical School**
James T. Elder
**University of New South Wales, NSW, on behalf of the Australia and New Zealand MS Genetics Consortium**
Bruce V. Taylor
**University of Ottawa**
Robert Dent, Ruth McPherson
**University of Oxford**
Rory Collins, Jonathan Emberson, Jason Torres
**University of Pennsylvania**
Brendan Keating, Joan O'Brien, Daniel J. Rader, Marylyn Ritchie, Dwight Stambolian
**University of Pittsburgh**
Adam Glassman, Erin E. Kershaw, Georgios Papachristou, David C Whitcomb
**University of Tasmania, on behalf of the Australia and New Zealand MS Genetics Consortium**
Nicholas B. Blackburn, Bennet J. McComish, Allan Motyer

**University of Texas Health Science Center at Houston**
Shervin Assassi, Maureen D. Mayes
**UT Southwestern Medical Center**
Jonathan Cohen
**Vanderbilt University Medical Center**
Eric D. Austin, Nancy J. Cox, Eric Gamazon
**Westmead Institute for Medical Research, on behalf of the Australia and New Zealand MS Genetics Consortium**
Grant P. Parnell
